# Robust detection of natural selection using a probabilistic model of tree imbalance

**DOI:** 10.1101/2021.05.12.443797

**Authors:** Enes Dilber, Jonathan Terhorst

## Abstract

Neutrality tests such as Tajima’s *D* (Tajima, 1989) and Fay and Wu’s *H* (Fay and Wu, 2000) are standard implements in the population genetics toolbox. One of their most common uses is to scan the genome for signals of natural selection. However, it is well understood that deviance measures like *D* and *H* are confounded by other evolutionary forces—in particular, population expansion—that may be unrelated to selection. Because they are not model-based, it is not clear how to deconfound these statistics in a principled way.

In this paper we derive new likelihood-based methods for detecting natural selection which are robust to confounding by fluctuations in effective population size. At the core of our method is a novel probabilistic model of tree imbalance, which generalizes Kingman’s coalescent to allow certain aberrant tree topologies to arise more frequently than is expected under neutrality. We derive a frequency spectrum-based estimator which can be used in place of *D*, and also extend to the case where genealogies are first estimated. We benchmark our methods on real and simulated data, and provide an open source software implementation.

## 1 Introduction

Understanding how species to adapt to their surroundings has been a defining challenge in biology for several centuries. One of the primary drivers of adaptation is, of course, natural selection. Recently, as genomic data has become much easier to obtain, significant efforts have been made to study natural selection using patterns of population genetic variation. In addition to advancing our general knowledge of evolution, this research has the potential to improve health and reduce disease by pinpointing the molecular basis for certain complex, adaptive phenotypes.

Because natural selection exerts a strong influence on the trajectory (frequency over time) of a selected allele, the ideal data for studying selection are time series of allele frequencies observed across many generations. Unfortunately, such data are rare except in laboratory settings. In order to study selection in natural populations, research has focused on devising methods for inferring selection from contemporaneous samples of polymorphism data. This is a challenging problem, because we have to make inferences about complex selection mechanisms using only a “snapshot” of genetic variation obtained at a single point in time. Theoretical models are essential in order to decipher these complicated signals in a principled way.

One way to reason about signals of natural selection is by considering its effect on genealogies. Relative to a neutral baseline, natural selection induces certain genealogical distortions. For example, a positively-selected variant sweeping towards fixation induces unbalanced, “star-like” genealogies, resulting in excesses of linkage disequilibrium and low- and high-frequency variants in the vicinity of the selected allele (Tajima, 1989; Fu and Li, 1993; Fay and Wu, 2000; Kim and Nielsen, 2004). Another form, balancing selection, produces genealogies which outwardly resemble those found in a structured population (Kaplan, Darden, and Hudson, 1988). These distortions are then manifested in terms of altered patterns of genetic variation. By fitting a statistical model of this process, we can learn about natural selection from polymorphism data.

### 1.1 Our contribution

In this article, we derive new procedures for detecting natural selection in genetic variation data. Our approach is based on a probabilistic model of genealogical imbalance which is designed to capture certain hallmark signals of selection described above. It generalizes Kingman’s ubiquitous coalescent process (Kingman, 1982a; Kingman, 1982b), and builds on earlier attempts in phylogenetics to model the process of speciation (Aldous, 1996; Blum and François, 2006). Although more principled and correct models of the coalescent process under selection have been studied previously (Krone and Neuhauser, 1997; Neuhauser and Krone, 1997), owing to their complexity, they are not widely used for inference. As we will see, ours is a simple approximation which retains much of tractability of neutral coalescent; the resulting estimators are fast, model-based, and easy to understand and implement. An important feature of our method it explicitly models variation in effective population size, leading to a “demographically corrected” neutrality test that has demonstrable advantages when population size indeed varies over time. Finally, because our method is based on a generative model of tree formation, it can be extended with little effort to cases where gene trees or ancestral recombination graphs have already been inferred, as is becoming increasingly common in population genetics (Kelleher et al., 2019a; Speidel et al., 2019).

### 1.2 Related work

We lack space to survey the full panoply of methods that have been developed to study natural selection using genomic data; see recent reviews by Vitti, Grossman, and Sabeti (2013) and Stern and Nielsen (2019). We focus here on two classes of methods for detecting natural selection which are most closely related to our proposed approach.

The first class is *frequency spectrum-based methods*, which operate on the principle that natural selection distorts equilibrium allele frequencies relative to what is observed under neutrality. The most widely used frequency spectrum-based statistic is Tajima’s *D* (Tajima, 1989):

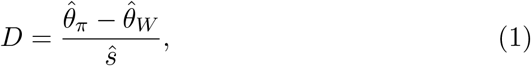

where 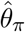 and 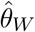 are, respectively, Tajima’s and Watterson’s estimators of the population-scaled mutation rate *θ*, and 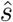 is an estimate of the standard deviation of their difference. Both estimators are unbiased for *θ* under neutrality, but have different biases for non-neutral evolution, such that 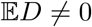 when examining allele frequencies obtained from a region that is under selection. Other related statistics include Fu and Li’s *D* (Fu and Li, 1993), and Fay and Wu’s *H* (Fay and Wu, 2000). A unifying interpretation of the various frequency spectrum-based statistics was given by Achaz (2009) who showed that each can be written as a certain weighted sum of entries of the SFS.

As suggested by (1), a common feature shared by all of the above-mentioned tests is that they are based on measures of deviance. That is, under neutrality each test statistic has zero mean, and larger magnitudes of the statistic suggest larger deviations from neutrality. However, beyond this general feature, interpretation of these measures can be subtle. For example, Tajima’s *D* is sensitive to deviations at all locations of the frequency spectrum, whereas Fay and Wu’s *H* only has power to detect a large excess of high frequency variants (Achaz, 2009). Negative values of *D* might indicate either directional selection or population growth, while positive *D* can alternatively indicate either balancing selection or population structure (Ferretti et al., 2017). More generally, deviance statistics based on the SFS are confounded by other evolutionary forces, in particular fluctuating historical effective population size, and there is not an obvious way to compensate for this^1^. Finally, because they operate using only marginal allele frequency information, these methods do not incorporate haplotype information or patterns of allele sharing, which can be a valuable auxiliary signal of natural selection.

A second group of methods for detecting selection can be described as *haplotype-based* methods. These are designed to exploit characteristic signatures of linkage disequilibrium that are deposited in the genome in the wake of a selective event (Maynard Smith and Haigh, 1974; Kaplan, Hudson, and Langley, 1989). Among the best-known of this class of methods are the so-called extended haplotype homozygosity (EHH) score (Sabeti et al., 2006), the integrated haplotype score (iHS; Voight et al., 2006), and the singleton density score (SDS; Field et al., 2016). Each of these scores is derived via population genetic and/or genealogical arguments about how variation is altered in the vicinity of a selected variant. For example, SDS is designed to detect regions of the genome where the terminal branches of the underlying genealogy are shorter than usual, as is expected under recent positive selection. However, although each of these statistics has been shown to work well in certain settings, ultimately these methods are heuristic, and not based on a concrete evolutionary model.

Given the profusion of *ad hoc* methods that have been proposed for detecting natural selection, it is natural to wonder why likelihood-based methods are not more common. The advantages of likelihood-based testing and estimation are well known (Neyman and Pearson, 1933; Lehmann and Casella, 2006). However, likelihood-based methods in population genetics are, in general, difficult: computing the likelihood of a sample of genomes, even under a simple neutral model, requires integrating over all of the possible ancestry scenarios that could have generated a given data set, a massive computational undertaking (Stern and Nielsen, 2019). Nevertheless, there has been some recent progress. Berg and Coop (2015) studied an approximate likelihood model for selection at a single locus, and very recently, a noteworthy contribution was made by Stern, Wilton, and Nielsen (2019), who propose an approximate full-likelihood method for inferring natural selection using recombining sequence data. Building on earlier work (Rasmussen et al., 2014), their method (approximately) integrates over the space of all possible allele genealogies and allele frequency trajectories for the selected allele.

Although these likelihood-based methods achieve state-of-the-art results, a potential downside is that they are computationally expensive. The method of Stern, Wilton, and Nielsen, for example, depends on obtaining a posterior sample of local trees from the program ARGweaver (Rasmussen et al., 2014), which can take many hours to generate even for moderate sample sizes. In practice, this makes it less likely that such methods would be employed in the exploratory phase of an analysis, as is routinely done with e.g. Tajima’s *D*. It seems that there is scope for a method that is easy to deploy while also mitigating some of the confounding issues described above.

## 2 Methods

Our starting point is the standard *n*-coalescent (Kingman, 1982a; King-man, 1982b) which is defined as a stochastic process on the set of par-titions of the set {1, …, *n*}. The process begins at time *t* = 0 in state 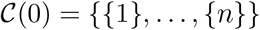. The instantaneous transition rate at time *t* is 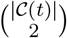 where 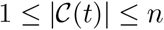 denotes the number of blocks in the partition remaining at time *t*. When a transition occurs, the new state is obtained by choosing two partition blocks uniformly at random and merging them. Thus, the number of partition blocks decreases monotonically over time, continuing until it reaches the absorbing state {{1, …, *n*}}. The trajectory of states 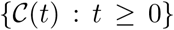 can be straightforwardly identified with a bifurcating tree on *n* leaves, with internal nodes occuring upon each block merger. For this reason, Kingman’s coalescent is often described as a distribution on binary trees.

An algorithm for drawing from Kingman’s coalescent follows directly from the above description. It is listed in the supplement (Algorithm S1) for completeness, though it is quite well-known. In this paper, we focus on an equivalent, but less common, method of sampling from Kingman’s coalescent, with the goal of obtaining a generalization which will prove useful for studying natural selection. This algorithm is shown in Algorithm 1. The main distinction is that it proceeds *forwards* in time (i.e., from past up to present), as opposed to Kingman’s original, retrospective process. That both the forwards- and backwards-in-time algorithms have the same distribution follows from e.g. Durrett (2008, Theorem 1.8).

### 2.1 The *β*-splitting family

We are motivated to consider Algorithm 1 because it can be generalized to produce alternative distributions on tree topologies. Observe that in line 5 of Algorithm 1, we could replace the uniform distribution with some other distribution on {1, …, *|B_i_|*−1}. For example, a distribution which, for each *|B_i_|*, placed mass 1/2 on 1 and |*B_i_*|−1, would produce unbalanced “caterpillar” trees with a large portion of external branches. Similarly, a distribution which placed all mass on (or near) |*B_i_|/*2 would produce trees which tend to be more “balanced” than is observed under Kingman’s coalescent. These two extremes produce the types of trees that we expect to form under certain types of natural selection, in particular directional and balancing selection.^2^

Such a model has been proposed by Aldous (1996), who studied probability distributions on random cladograms (topological trees with no branch length information). Aldous defined a one-parameter family of distributions which he called the *β-splitting model*^3^. In this model, a clade of size *n* is randomly split into subclades of sizes {*i,n−i*}, where now *i* is distributed according to a symmetric beta-binomial distribution with shape parameter *β*, conditioned on *i* ∉ {0,*n*}. Concretely, this distribution is given by

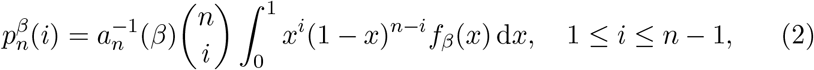

where

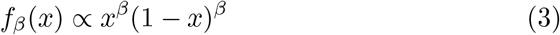

is the symmetric beta density with shape parameter *β*, and

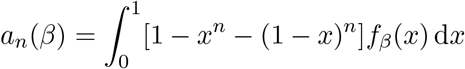

is the normalizing constant. Integrating out *x* in (2), one obtains

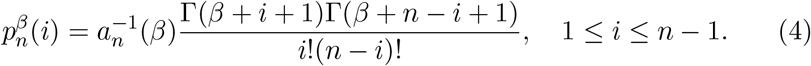

The beta density (3) is integrable for *β* > −1 (note that Aldous’ parameterization differs by 1 from the usual convention.) However, up to normalization, (4) defines a valid probability distribution whenever min(*β* + *i* + 1*, β* + *n* − 1 + 1) > 0; that is, for *β* > −2. For *β* = 0, *p_n_*(*i*; *β*) ∝ 1 and the distribution reduces to Kingman’s coalescent. By examining the ratio

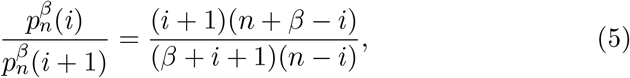

we see that letting *β* → ∞ causes 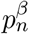 to place most of its mass near *n/*2, leading trees which are more “balanced” than under the usual coalescent. If *β* → −2, ratio in (5) diverges for *i* ∈ {1, *n* − 1}, so 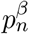 places mass on *i* ∈ {1, *n* − 1}, resulting in maximally unbalanced splits and a “caterpillar” tree.

The reader may wonder why the beta-binomial distribution was chosen, when we could conceivably have used any distribution on {1, …, *n* − 1}. The symmetric beta-binomial is attractive due to parsimony (it adds only one extra parameter), and because it preserves some desirable properties of tree distributions such as exchangeability. Also, its usage has precedent in the related field of phylogenetics, where it has been proposed as a model for speciation (Blum and François, 2006). Other authors have recently studied further generalizations of this process to the case where the shape parameters are not symmetric (Sainudiin and Véber, 2016). A disadvantage of this model is that, in contrast to Kingman’s coalescent, the forward-splitting process does not seem have any evolutionary interpretation (Aldous, 1996). We choose to view it empirically as a useful tool for studying natural selection using coalescent-based methods.

### 2.2 Expected site frequency spectrum

Given a sample of *n* individuals, the expected site frequency spectrum (ESFS) is the distribution of the number of individuals *i* ∈ {1, 2, …, *n* − 1} bearing the derived allele at a randomly selected segregating site. (We assume that the identity of the ancestral allele is known.) In this section we show how to determine the ESFS under the *β*-splitting model.

We denote the ESFS by 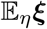, where the site frequency spectrum ***ξ*** ∈ Δ^*n*−1^ is the sample version of ESFS, i.e. a vector whose *i*^th^ entry denotes the proportion of segregating sites where *i* members of the sample bear the derived allelle. Here Δ^*n*−1^ denotes the (*n* − 1)-dimensional probability simplex, i.e. the set of all numbers *x*_1_*, …, x_n_* ≥ 0 such that *x*_1_ + · · · + *x_n_* = 1. The expectation is taken with respect to genealogies generated under a given evolutionary model *η*. Although *η* could in principle be quite general, efficient methods for computing 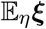 are only known when *η* describes neutral evolution under either constant or variable effective population size. Therefore, from this point on we take *η* to represent a function representing the historical size of the population.

Under an “infinite sites” model with low rates of mutation, Griffiths and Tavaré (1998) have shown the following key result:

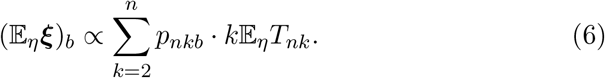

In the preceding display, 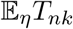 is the average amount of time (under the evolutionary model *η*) during which there are *k* lineages ancestral to a sample of size *n*, and *p_nkb_* the probability that a branch at level *k* in an *n*-coalescent tree has *b* sampled descendants in the present.

In Kingman’s coalescent,

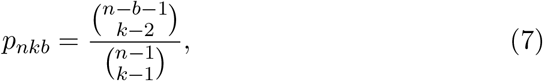

which can be derived by a combinatorial “stars-and-bars” argument (Durrett, 2008). If the effective population size is constant, then 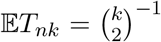 from which follows the well known result that 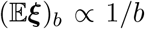 for Kingman’s coalescent. If population size varies through time according to some size history function *η*(*t*), then a simple expression for 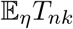 no longer exists, but Polanski and Kimmel (2003) have shown that it may be computed as a certain linear transformation of the vector of first coalescent times _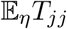_, *j* = 2, …, *n*. We return to this fact below.

Although Kingman’s coalescent and its generalization to variable effective population size are the two best-known applications of Griffiths and Tavare’s formula (6), in fact their argument holds more generally for any distribution on trees, assuming (crucially) that the branch lengths and topology of those trees are independent. Since this is true for the *β*-splitting model defined above, we can use a generalization of (6) to derive its expected SFS.

Let _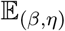_ denote expectation with respect to trees generated under the *β*-splitting model. Since the *β*-splitting model alters tree topology only, we have

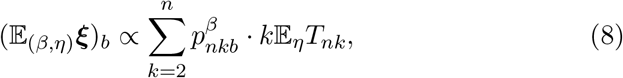

where the vector 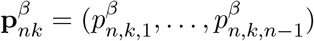 has the same interpretation as above. In the next two subsections, we show how to compute the “topological” 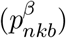 and “branch length” 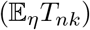 components of this formula.

#### 2.2.1 Dynamic programming algorithm for 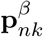

A simple expression like (7) does not seem to exist when *β* ≠ 0. Instead, we derive a dynamic programming algorithm for calculating the combinatorial factors 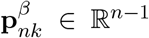, *k* = 2, …, *n* defined in the preceding section. The method applies to any forward-splitting model and includes *β*-splitting as a special case.

Define 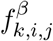 to be the probability that a size-*i* block at level *k* splits into blocks of size *j* and *i − j*. From the preceding section, we know that under Kingman’s coalescent,

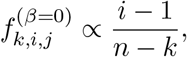

and for the general *β*-splitting model,

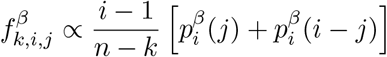

where 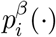 was defined in equation (4).

For each level *k* let S*^k^* ∈ ℤ*^n^* be a row vector such that 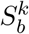is number of nodes at level *k* which subtend *b* = 1, …, *n* leaves at the bottom of the coalescent tree. Also let **e**_1_, …, **e**_*n*_ ∈ ℝ^*n*^ be the standard basis (row-)vectors. Under the forward-splitting model described above, the sequence **S**^1^, **S**^2^, …, **S**^*n*^ forms a Markov chain, with transition probabilities

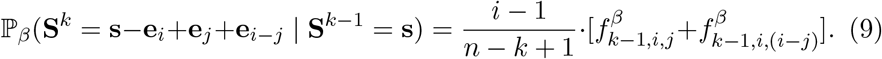

The starting state of the Markov chain is **S**^1^ = (0, 0*, …*, 1) = **e**_*n*_. Focusing on an individual entry 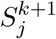 and summing over all possible events that would cause it to *increase*, we obtain

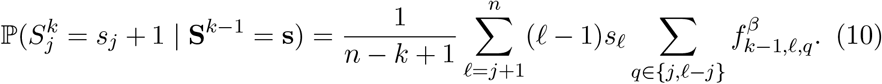

Similarly, a *decrease* can occur only if a size-*j* block was chosen to split in the preceding level:

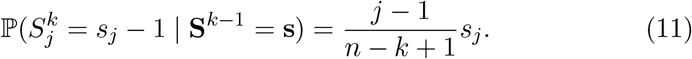

In matrix notation, (10) and (11) combine to yield

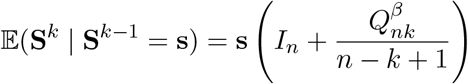

where *I_n_* ∈ ℝ^*n*×*n*^ is the identity matrix, and

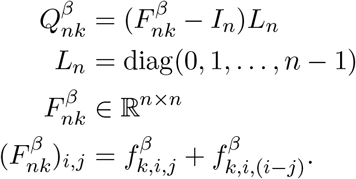

Hence,

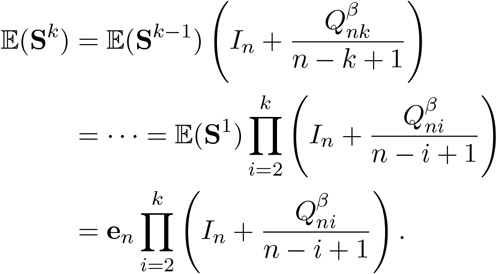

Finally,

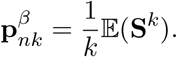

#### 2.2.2 Computing the expected branch lengths

Next we discuss how to compute the other necessary quantity 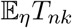 in equation (8). Let **T***_n_* = (*T_n,_*_2_*, T_n,_*_3_, …, *T_n,n_*) be the vector of these times. Polanski, Bobrowski, and Kimmel (2003) have shown the following relationship for a general size history function *η*:

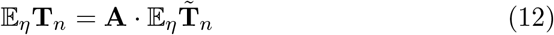

where 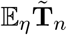 is the vector of first coalescent times,

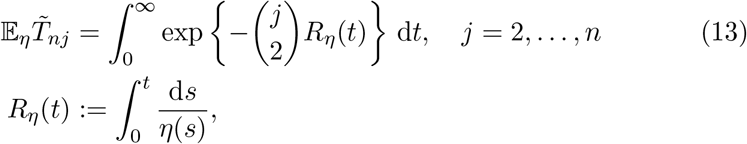

and A*_n_* ∈ ℝ^(*n−*1)*×*(*n−*1)^ has entries

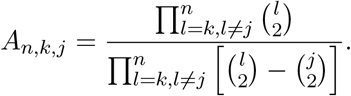

As in the preceding section, this result holds for any tree distribution in which branch lengths and topology are independent, so it can be applied to our model.

Readers who are familiar with this area may notice that, for Kingman’s coalescent, the expected SFS is typically not calculated via equation (8). Instead, by another result of Polanski and Kimmel (2003), interchanging the order of summations in equations (8) and (12) allows the (unnormalized) ESFS to be expressed as a linear transformation of 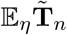. Unfortunately, this trick does not lead to simplifications in our more general model, so we first compute the expected intercoalescence times and then plug them into (8). For large *n*, the matrix-vector product (12) is numerically unstable, so we use a high precision numerical library to evaluate the integral (13) and then (12). This approach is less efficient than using hardware floating point operations, but it only needs to be performed once per given demography, so it is suitable for genomewide analysis.

### 2.3 Estimating *β*

Given the probabilistic model defined above, how can we estimate it in order to infer *β*? In this section, we propose two methods depending on the type of data that are available.

#### 2.3.1 From the SFS

To perform inference using the SFS we rely on the so-called *Poisson random field* (PRF) approximation (Sawyer and Hartl, 1992), which assumes the coalescent tree at every segregating site is independent of all others.

Assuming also that mutations are rare—formally, that *θ* → 0, as is reasonable for humans and many other species—then we may approximate the mutation process on a coalescent tree by a Poisson process.

Given an empirical frequency spectrum *ϕ* ∈ ℤ^*n*−1^, where *ϕ_i_* is the number of segregating sites where *i* copies of the derived allele were observed, the PRF log-likelihood is

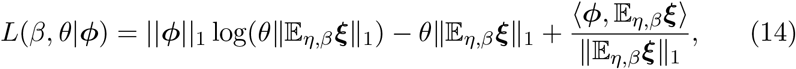

where the ESFS 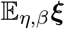 is calculated using the procedure derived in Section 2.2. If the mutation rate *θ* is not known, then the maximum likelihood estimate can be shown to equal

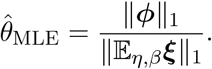

Substituting this back into (14), and setting p = *ϕ*/||*ϕ*|| _1_, _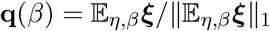_, we obtain that the profile likelihood

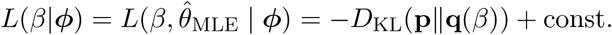

In order words, maximizing the likelihood is equivalent to minimizing the KL divergence between the categorical distributions **p** and **q**(*β*) (Bhaskar, Wang, and Song, 2015).

#### 2.3.2 From inferred trees

The ESFS is obtained by integrating over all possible genealogies at a given site, and then fit to data by assuming independence between sites. An alternative strategy is try to estimate those genealogies, and then do inference conditioned on them. Recently in population genetics, there have been methodological breakthroughs that enable the estimation of ancestral recombination graphs using large numbers of genomes (Kelleher et al., 2019a; Speidel et al., 2019). In the future, as algorithms and computational capabilities continue to improve, this may become the dominant mode of population genetic analysis. We therefore explored extensions of our methods to the case where genealogies are estimated instead of integrated out.

Because of the probabilistic nature of our model, it is easy to extend it to the case where the genealogy is observed instead of latent. Moreover, estimating *β* conditional on a collection of inferred genealogies simplifies the problem considerably. If we assume a bifurcating tree, the sizes of children nodes can be modeled by the beta-binomial distribution as previously described. Just like the preceding section, we proceed level by level in the (now observed) genealogy. At each level *k* = 2, …, *n* of the tree, let the size of the internal node which splits into two child nodes be denoted *B_k_*, and the sizes of its child nodes *c_k_* and *B_k_ − c_k_*. We model the probability of an the observed tree 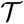 as

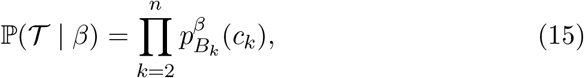

with 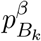 defined as in (4), so that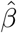 obtained by numerical optimization.

##### Weighted likelihood

When experimenting with this method, we observed a small but consistent performance improvement by reweighting the likelihood (15):

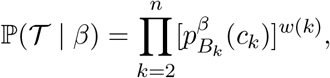

where *w*(*k*) is a weighting function. For detecting directional selection, we found that setting the weights proportional to the size of the internal node, *w*(*k*) = *B_k_*, worked well. For detecting balancing selection, we found that it helped to weight the various terms by total amount of branch length at their respective level in the tree: *w*(*k*) = *kt_k_*, where *t_k_* is the amount of branch length at level *k* in the tree (see Section 3.1.) Using weights improved the method’s performance of detecting the imbalance of the tree. The effect of the different weighting methods is shown in Figures S4 and S5. The gain was around 0.01–0.04 AUC in each scenario.

##### Related tree imbalance statistic

The Colless statistic (Mooers and Heard, 1997) is a measure of the imbalance of a binary tree, defined as

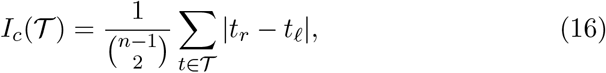

where the summation is over all internal nodes *t* of the tree, and *t_r_, t_ℓ_* are the sizes of the two child nodes descending from *t*. We used the Colless statistic as a baseline for comparing the performance of our 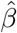 statistic when fitted to inferred trees. The exact relationship between 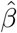 and 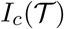 is somewhat opaque, but in general we can note that *I_c_* is maximized for a caterpillar tree, and is zero for a perfectly balanced tree with an even number of leaves.

Hence it is negatively associated with *β*-splitting parameter. In the next section, we compare the ability of these two measures to detect signals of selection.

##### Polytomies

In practice, we found that current tree inference softwares often generate multifurcating trees. Since our method assumes a bifurcating tree, we first resolved these polytomies by arbitrarily breaking them into sequences of bifurcation events. Of course, polytomies could well represent additional selection signal. Our current implementation ignores this, but we discuss potential extensions in Section 4.

### 2.4 Alternative parameterization

We conclude this section with a note on implementation. When fitting our model to data, we observed that the parameterization (4) exhibited some numerical instability when performing gradient-based optimization. The problem arises when computing the normalizing constant for the range −2 < *β* < − 1 which, as mentioned in Section 2.1, can no longer be interpreted as a draw from a conditioned beta-binomial distribution. To work around this, we restrict *β* > − 1 and then perform a log transformation. Specifically, in all of the results reported below, the following alternative definition of the symmetric beta-binomial distribution is used:

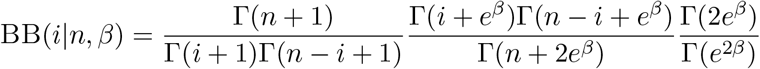

Then we restricted *i* to be in {1, 2, …, *n* − 1};

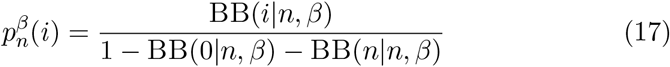

where *i* ∈ {0, 1, …, *n* − 1}, *n* ∈ ℕ^+^ and *β* ∈ ℝ. The transformed distribution has the following properties: when *β* = 0, this becomes a uniform distribution so the model recovers the usual Kingman’s Coalescent. When *β* → −∞, most of the weights of the distribution will be at the tails, so corresponding tree will be similar to a caterpillar tree. And when *β* → ∞, the weights will be accumulated around the center and lead to a balanced tree.

### 2.5 Data analysis pipeline

A description of the pipeline used to analyze data and run our methods is contained in the supplement (Section S1).

## 3 Results

In this section, we study various characteristics of the methods we derived in Section 2 using simulations, before concluding with applications to real data.

### 3.1 Topological variance analysis

Recently Ferretti et al. (2017) gave an interpretation of several frequency spectrum-based neutrality tests in terms of tree imbalance. In this section we study our model using some of their results. This helps clarify the connection between some existing neutrality tests and our work.

Following Ferretti et al., we define *d_k_* to be the size (number of leaf nodes subtended by) a randomly selected lineage at level *k* in a genealogy. Averaged over genealogies under the *β*-splitting model, we have

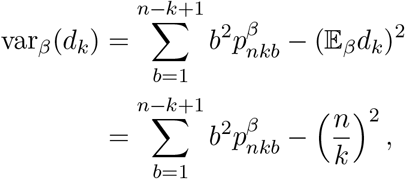

where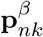 was defined in Section 2.2.1, and the second inequality holds because 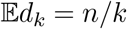 under any leaf-exchangeable tree distribution. Computing var*_β_* (*d_k_*) in closed form for our model is challenging due to the fact that 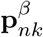 is recursively defined. Here we focus on a few special cases where we can derive a precise answer, and study the general relationship using simulations.

For *β* → −2, corresponding to the caterpillar tree, it is easy to show that

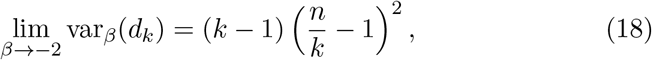

as already noted by Ferreti et al. Also, for Kingman’s coalescent, *β* = 0,

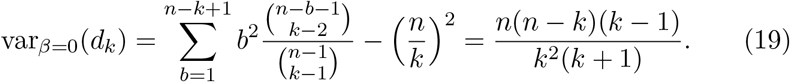

For *β* → ∞, we were unable to derive a closed-form expression for lim_*β*→∞_ var_*β*_ (*d_k_*). However, Ferretti et al. showed that the dominant contribution to topological variance comes from level *k* = 2, for which

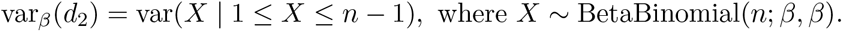

If *n* and *β* are both large, the condition 1 ≤ *X* ≤ *n* − 1 has probability near one and can be ignored. Using the variance formula for the beta-binomial distribution, we have

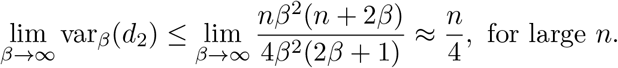

We further define

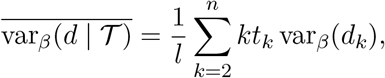

which is the topological variance of a given genealogy, weighted by the relative proportion of branch length at each level (see equation (4) in Ferretti et al.). Substituting *t_k_* and *l* by their expected values in equations (18) and (19), as *n* → ∞,

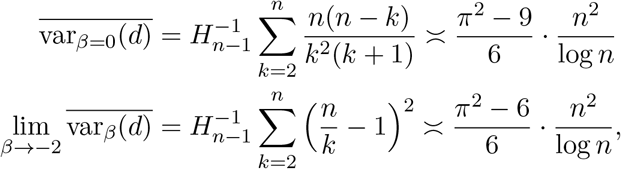

where *H_n_* is the *n*th harmonic number.

Now let *T* be a neutrality test statistic (for example, Tajima’s *D* or Fay and Wu’s *H*). Since the parameter *β* only affects tree topology, we obtain from formula (17) of Ferretti et al.,

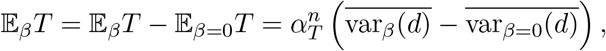

where *α_T_* (*n*) is a test-specific constant which depends on *n*, and for simplicity we ignored the normalization term *f*_Ω_(*θl*).

To show an example of how the topological variance affects neutrality tests such as Tajima’s *D*, we simulated genealogies under various settings of *β*, assuming constant population size with no recombination (Figure 1). The box plots are empirical distributions of two neutrality tests (Tajima’s *D* and Fay and Wu’s *H*) for various settings of *β* ∈ [− 2, ∞). The dashed red lines represent the limiting values predicted by the calculations shown above. The figure shows how different values of these statistics can be interpreted in terms of *β*, and vice versa. We see, for example, that *D* and *H* appear to be more sensitive to *β <* 0, in the sense that their distribution at *β* = 0 nearer the *β* → ∞ limit than the *β* → −2 limit.

**Figure 1:**
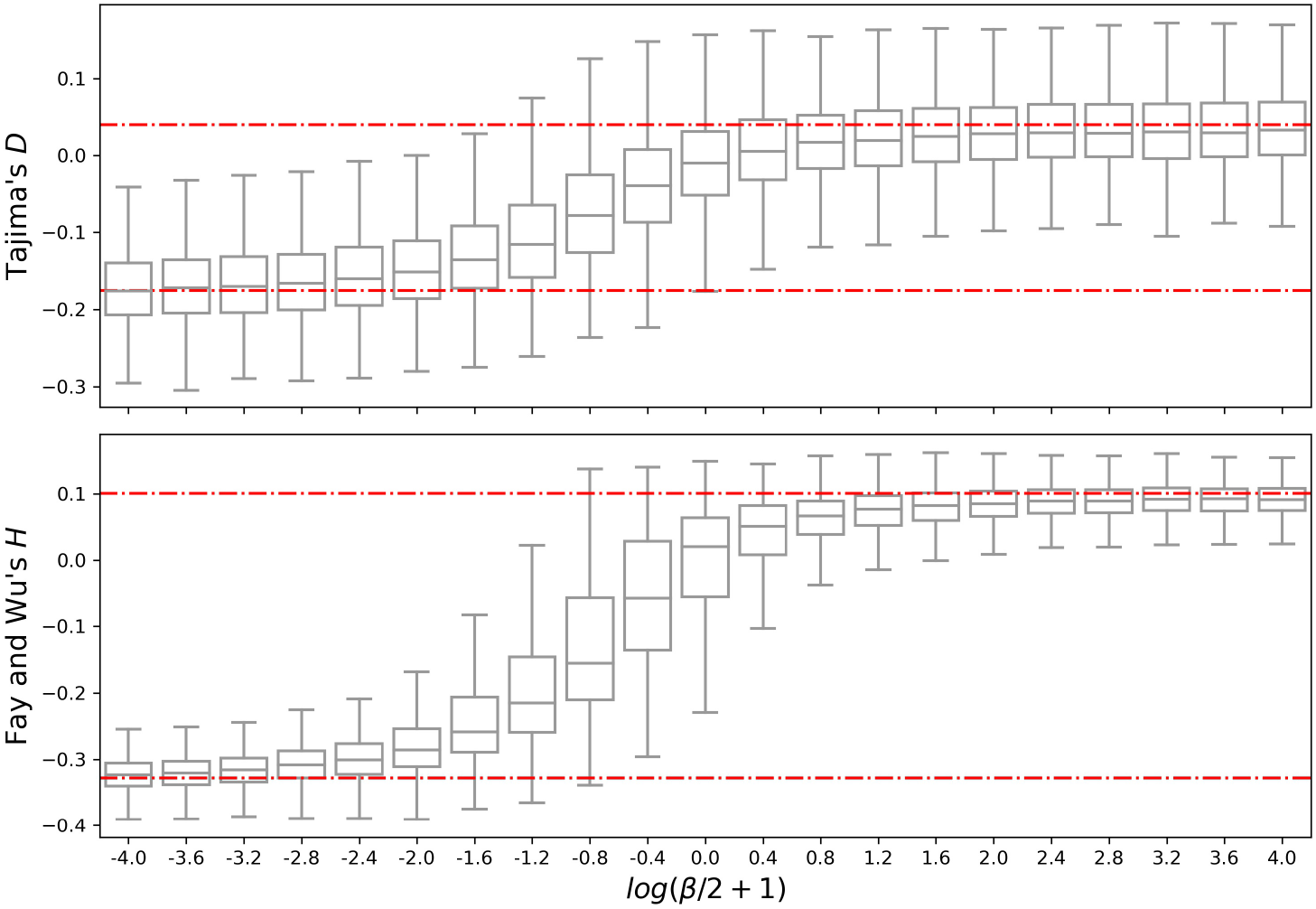
Empirical distributions of Tajima’s *D* and Fay and Wu’s *H* under different tree topologies. Going from left to right, tree structure goes from caterpillar to balanced. Red lines represent the averaged limiting cases.

### 3.2 Simulated data

To benchmark our methods on simulated data, we studied their ability to classify simulated genomic regions as being either neutral or under some form of selection. The receiver operating characteristic (ROC) curve, and associated area under curve (AUC) statistic, are standard ways to measure the performance of a classifier. For each experiment described below, we generated data under two different models, and then plotted ROC curves for each method. The two possible models are printed at the top of each ROC curve. The legend lists each method that was compared, along with its AUC score.

The classification procedures derived from our methods are denoted `btree` and `bsfs`. The `bsfs` results were obtained by maximizing (14) over *β* with respect to the observed frequency spectrum. `btree` is the tree-sequence based estimate, obtained by maximizing the conditional likelihood defined in (15) over *β* conditional on a given tree. As a baseline, we also compared our method to Colless’ statistic (see Section 2.3.2) and Tajima’s *D*. Finally, ROC curves were computed by thresholding the empirical null distributions of each test statistic. We also use these neutrally evolved simulations to infer population size histories (*η*(*t*)) that we use for `bsfs`. Our simulation process is explained in detail in Section S1.1.

#### 3.2.1 Directional selection

We simulated a single population with constant population size *N* = 2 × 10^4^. The simulated region was 10^5^ base pairs, with recombination and mutation rates of 1.25 × 10^−8^ and 2.5 × 10^−8^ per base pair per generation, respectively. When each simulation terminated, we randomly sampled *n* = 50 haploid genomes and computed the relevant test statistics. We introduced a beneficial mutation 250 generations prior to present into the middle of the region, and we restarted the simulation if the mutation is lost or fixed. Following Stern, Wilton, and Nielsen (2019), we varied two parameters; selection co-efficient *s* ∈ {.001,.003,.01,.02} and allele frequency *F* ∈ {0.25, 0.5, 0.75} of mutation when the simulation terminated. Genic selection was assumed, i.e. the relative fitnesses of the wild-type homozygotes, heterzygotes, and derived homozygotes were 1, 1 + *s/*2, and 1 + *s*, respectively.

Figure 2 displays results for each of the methods. In general, we observed that tree-sequence based methods are better at detecting strong selection compared to SFS-based methods. This is expected, because a recent hard sweep leaves a signal of elevated linkage disequilibrium that is invisible in the frequency spectrum (Kaplan, Hudson, and Langley, 1989). In particular, the `btree` method achieves at least 0.8 AUC for *s* ≥ 0.003 and *F* ≥ 0.5. The performance Colless’ statistic and `btree` are similar. `btree` has significantly higher AUC (*p* =.014, Wilcoxon signed rank test), but the overall gain is small (mean ΔAUC = 0.0034). Among the SFS-based statistics, our method (`bsfs`) achieved significantly higher AUC scores (*p* =.0014, Wilcoxon signed rank test) than Tajima’s *D*, and the average gain is notable (mean ΔAUC = 0.049).

**Figure 2:**
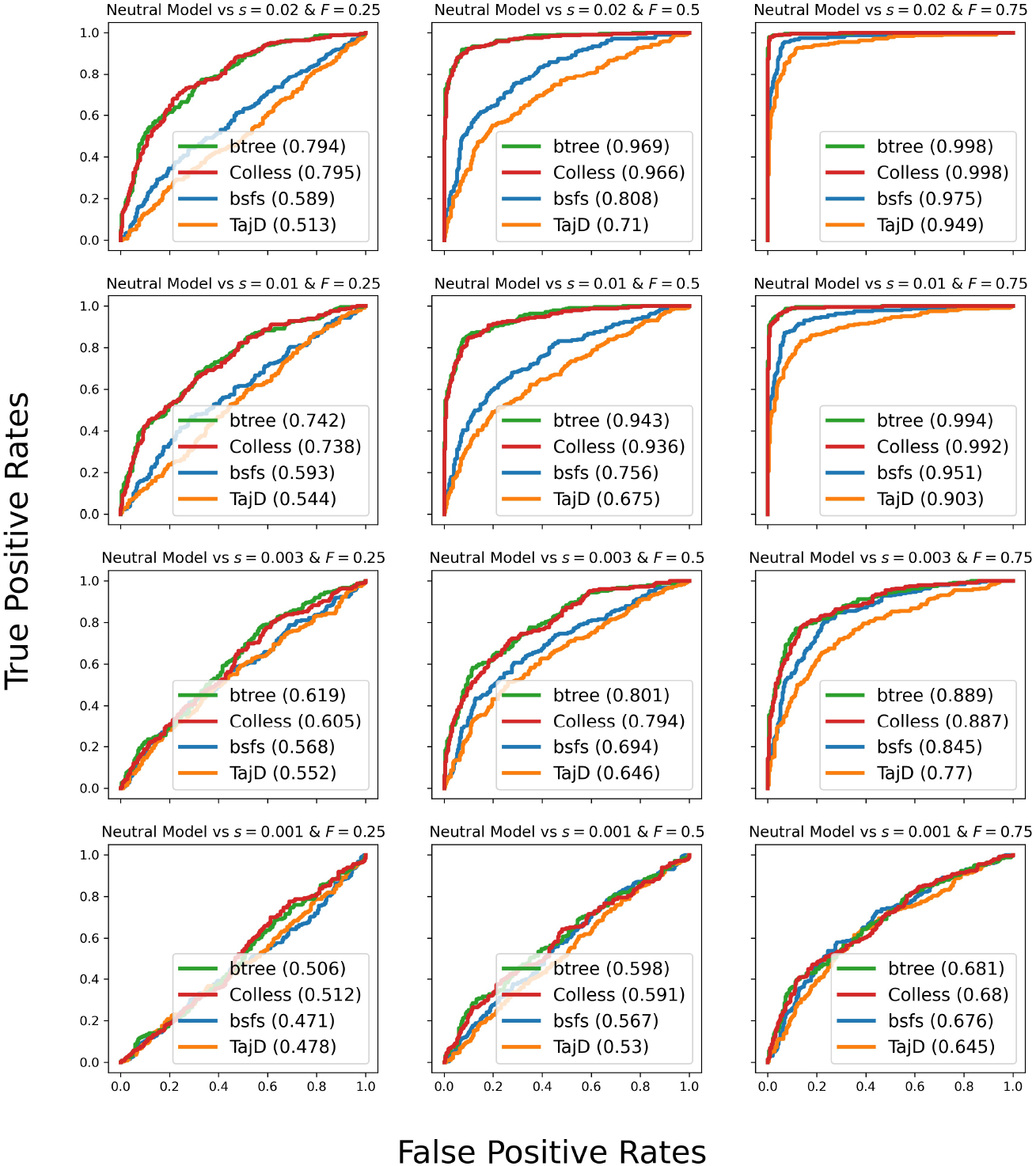
ROC curves for positive genic selection. *s* represents selective advantage of the mutation and *F* represents allele frequency of the mutation in the sample.

#### 3.2.2 Balancing selection

Next we studied our methods’ ability to detect long-term balancing selection. Since this type of selection acts on a longer time scale than directional selection (Charlesworth, 2006), it is necessary to forward simulate for many more generations. To speed up the simulations, we reduced the population size by a factor of 10 to *N* = 2 × 10^3^, and increased the mutation and recombination rates to 1.25 × 10^−7^ and 2.5 × 10^−7^. The simulated region was 2500 base pairs. When each simulation terminated we randomly sampled *n* = 250 haploid genomes and computed the relevant test statistics using them. Heterozygously advantageous mutations were introduced at constant rate throughout the simulation. We varied two parameters: *t*_0_ ∈ {2 × 10^3^, 3 × 10^3^, 4 × 10^3^, 5 × 10^3^} which represents the number of generations before present when beneficial mutations began, and selection coefficient *s* ∈ {.0004,.0008,.002}. The dominance parameter was set to *h* = 25 in all cases. Thus the fitnesses of the homo- and heterozygote were ≈ 1 and *s* · *h* ∈ {.01,.02,.05}, respectively.

Figure 3 contains ROC curves along with the AUC values in parenthesis for each of the methods. For balancing selection `btree` again outperforms Colless’ statistic, but the difference is subtle (mean ΔAUC = 0.0026) and not significant (*p* = 0.31, Wilcoxon signed rank test). In contrast to the case of directional selection, SFS-based statistics did better than tree-based statistics in this example. Among the SFS-based statistics, our method (`bsfs`) achieved significantly higher AUC scores (*p* = 0.0011, Wilcoxon signed rank test) than Tajima’s *D* with a mean ΔAUC = 0.024. We performed some additional analysis to better understand why SFS-based statistics are better than the tree-based ones for detecting balancing selection. We found that long branches near the root of the tree that occur in genealogies under long-term balancing selection have a pronounced impact on the SFS, but do not affect the topology of inferred trees.

**Figure 3:**
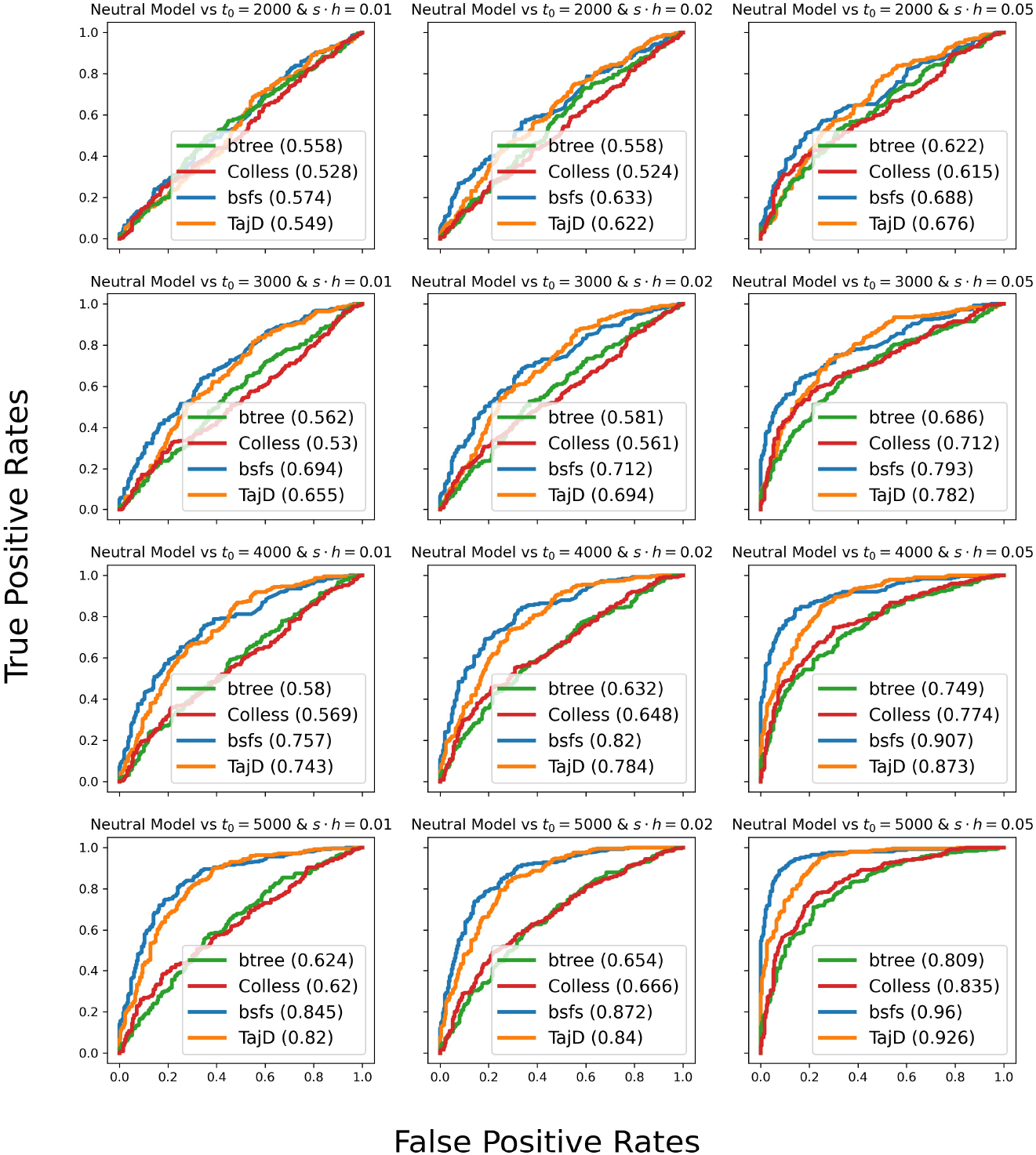
ROC curves for advantageous heterozygote mutation simulations. *s* represents selective advantage of the mutation, *h* is the dominance factor. *t*_0_ represents how many generations ago the advantageous mutations were introduced into the sample.

#### 3.2.3 Effect of variable population size

It is well known that, when used to detect natural selection, Tajima’s *D* is confounded by population structure and changes in effective population size (Stajich and Hahn, 2005; Biswas and Akey, 2006). In the single-population case, one interpretation of this phenomenon is that *D* measures both topological and branch length distortions compared to the neutral coalescent (Ferretti et al., 2017), and population size changes also distort branch lengths. In contrast, our SFS-based estimator is designed to detect topological changes only, and it can be modified to take into account population size history (Section 2.2).

We compared the ability of *D* and bsfs to detect directional selection under four scenarios:

- Constant population size under neutrality;
- Exponential growth under neutrality;
- Constant population size with directional selection;
- Exponential growth directional selection.

For the selective scenarios, we introduced a single mutation 250 generations before present to the middle of the 10^5^ base pair region, restarting the simulation if the mutation was lost or fixed. The sample size was *n* = 250 haploids. The recombination and mutation rates were again 1.25 × 10^−7^ and 2.5 × 10^−7^. For the `bsfs` method, we first estimated the underlying population size history *η*(*t*) using 25Mb of neutral data simulated under the corresponding demography. Other varying parameters for the experiments can be seen at Table S1. In the table, *N_e_*(0) is the population size at the time simulation starts, *g* is the growth rate of exponential growth, *s* is the selective coefficient of the beneficial mutation and *h* is the dominance parameter.

In Figure 4a, our method has higher AUC than *D* for distinguishing a neutral model from selection for both constant population size and exponential growth (left and center panels). To illustrate the pitfalls of using *D* without correcting for demography, we also considered a third scenario (right-most panel) in which there is *no* selection; the only difference between the two models is that one of them underwent exponential growth, while effective population size in the other was constant. In this plot, a “true positive” signifies that the constant-sized model is rejected in favor of the exponential growth model when the latter model generated the data, and similarly for a false positive. As expected, the plot shows that *D* has high power to detect exponential growth—however, if the analyst were unaware that the population had experienced growth, then this could wrongly be interpreted as evidence for selection. In contrast, after adjusting the expected frequency spectrum to compensate for this effect, our estimator does no better than a coin-toss (AUC ≈ 0.5) at distinguishing between the two régimes.

**Figure 4:**
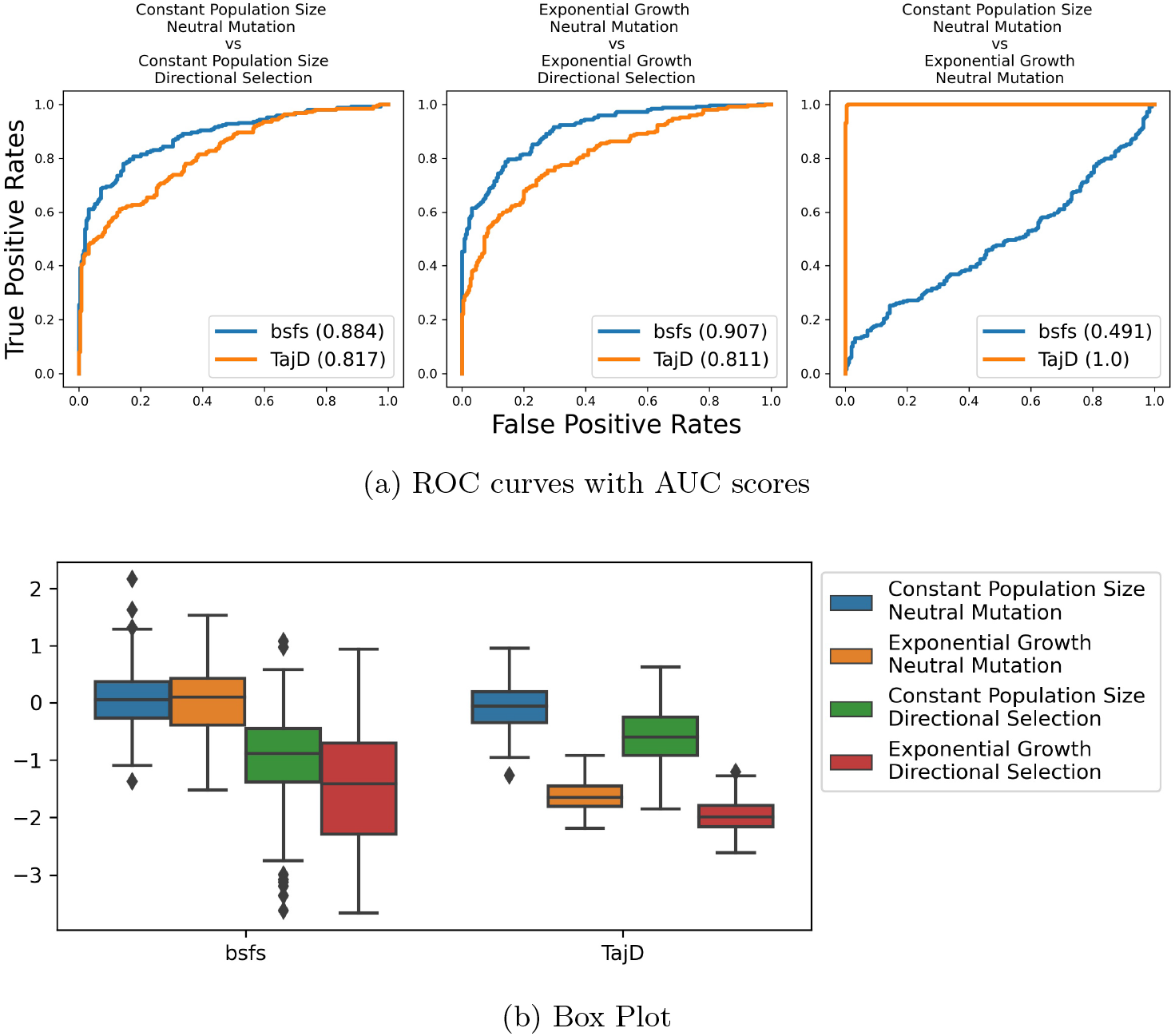
Directional selection under constant population or exponentially growing size histories. `bsfs` is our SFS based method and `TajD` is Tajima’s *D*. (a) `bsfs` performs better for detecting true signals in the first two figures. In the third figure, *D* is picking up a false positive signal with respect to detecting selection. (b) Under neutrality, `bsfs` has a zero centered empirical distribution regardless of the true population size history, whereas the distribution of *D* is shifted.

Another way to see this result is in Figure 4b, which shows the empirical distributions of *D* and 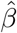 obtained from bsfs. After correcting for demography, the two neutral simulations (orange and blue) have roughly the same empirical distribution using our method, even though they are generated under quite different growth models. In contrast, the distribution of *D* under neutral exponential growth closely matches that of directional selection under exponential growth, and is very different from the distribution under neutrality and constant population size.

We repeated this experiment under simulated balancing selection. We again simulated four different scenarios:

1. Constant size with no advantageous mutation;
2. Exponential growth with no advantageous mutation;
3. Constant size with heterozygote advantage and;
4. Exponential growth with heterozygote advantage.

For the exponential growth scenarios, the growth began 250 generations ago. Detailed settings for each type of simulation are shown in Table S2. Results were similar to the directional selection experiment. In Figure 5b, we see that selection and growth “cancel out” in Tajima’s *D*: it has a similar distribution under exponential growth and balancing selection as under neutrality with constant size. In contrast, the null distribution of `bsfs` is invariant after correcting for demography.

**Figure 5:**
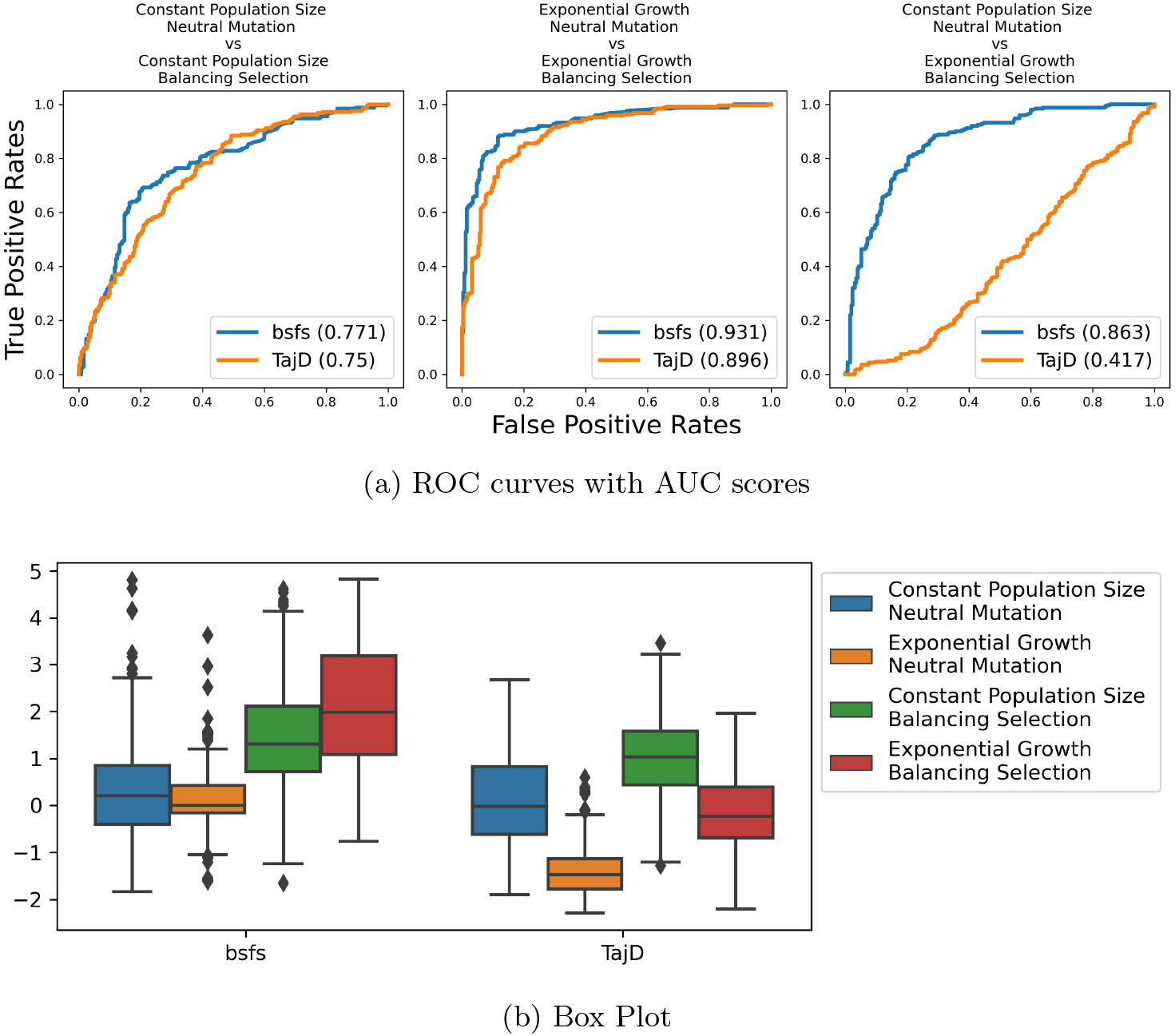
Balancing selection under constant population or exponentially growing size histories. (a) `bsfs` performs better for detecting true signals in the first two figures. In the third figure Tajima’s *D* fails to detect selection. (b) Under neutrality, bsfs has a zero-centered empirical distribution and balancing selection shifts the distribution upward. Balancing selection shifts *D* to positive values but exponential growth pulls it downward.

### 3.3 Real data analysis

We applied our models to data from the 1000 Genomes Project (The 1000 Genomes Project Consortium, 2015), using tree sequences that were inferred by Kelleher et al. (2019b). To understand how our model works compared to other known statistics, we focused on 7 regions which are known to experience selection: *LCT* in chromosome 2, *SLC45A2* in chromosome 5, *HERC2* in chromosome 15 for European populations; *SLC44A5* in chromosome 1,

*EDAR* in chromosome 2, *ADH1* in chromosome 4 for East Asian populations; *MHC* in chromosome 6 for all populations. For *LCT*, *SLC45A2*, *HERC2*, *SLC44A5*, *EDAR*, *ADH1* we used the `btree` statistic to investigate directional selection since it is sensitive to linkage disequilibrium. For *MHC* we used `bsfs` since our simulation results show that our frequency spectrum-based methods are better at detecting long-term balancing selection. We performed one-sided testing: for directional selection, *p*-values were calculated by *p^−^*, and for balancing selection by *p*^+^ (cf. eqn. 21).

#### 3.3.1 Directional selection

Lactose is the principle sugar in milk. Like other mammals, humans historically lost the intestinal enzyme lactase after infancy, and with it the ability to digest milk. But between 5,000 to 10,000 years ago, a genetic mutation arose that confers lactase persistence in adults. Today it is found in a majority of the adult populations of Northern and Central Europe. The location of this mutation in the gene *LCT* displays one of the strongest signals of directional selection in the human genome (Bersaglieri et al., 2004).

In Figure 6a, as expected we have a very small *p*-value for the European populations around *LCT*. This indicates our estimated *β*-splitting parameters are negative, as expected for strong directional selection (Section 2.1). Specifically, Utah Residents with Northern and Western European Ancestry (CEU), British in England and Scotland (GBR) and Finnish in Finland (FIN) have significantly negative 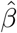. Southern European populations such as Toscani in Italia (TSI) and Iberian Population in Spain (IBS) also show evidence of selection, though the signal is weaker, reflecting the fact that the strength of selection may be lower in these populations (Gerbault et al., 2011).

**Figure 6:**
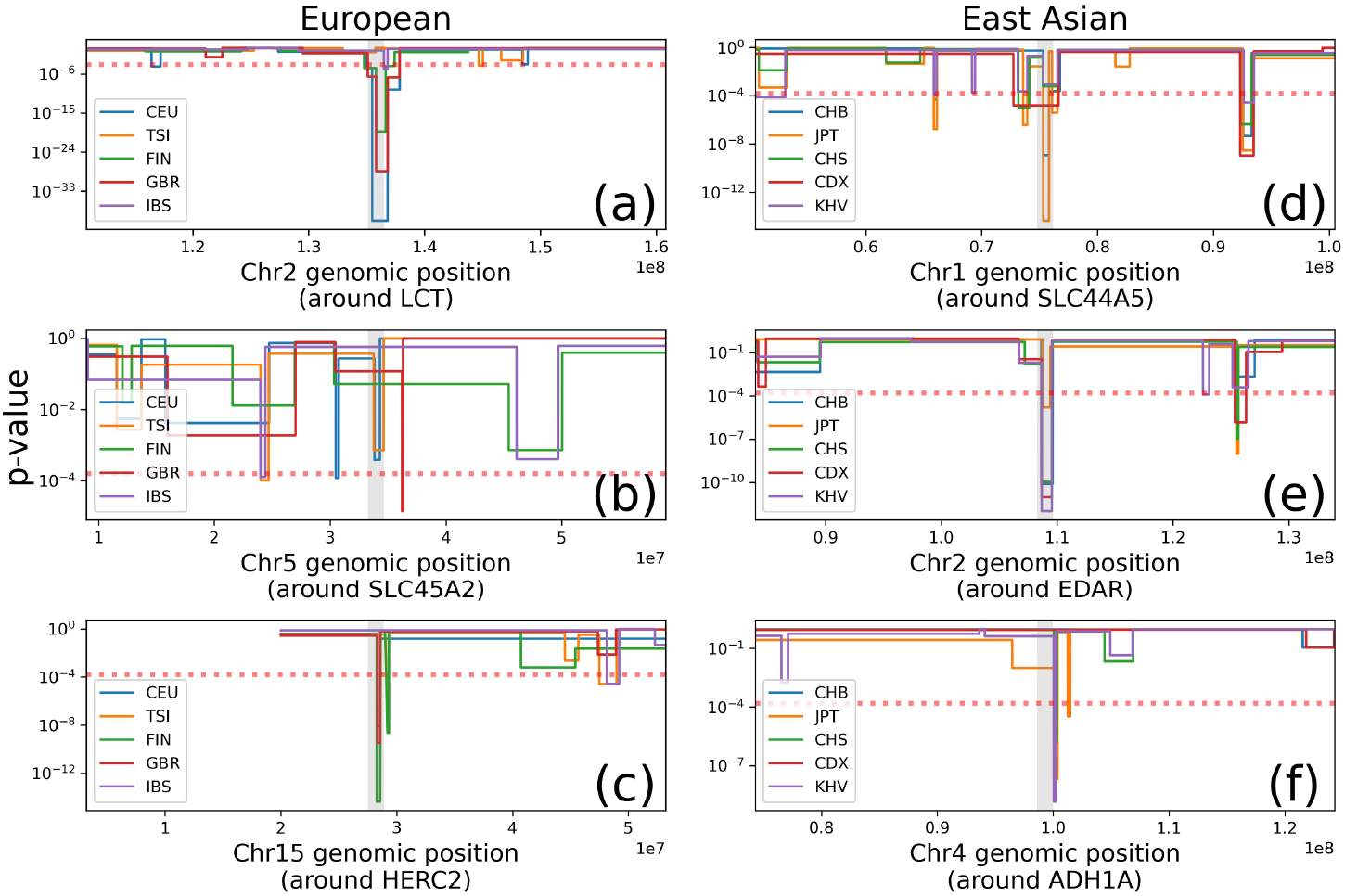
Results of directional selection *p*-value scan for 1000 Genomes Project using median centered `btree` (Section S1.2.1). The Bonferonni-corrected significance level is 1.6 × 10^−4^ (Red dashed line). Significant populations for each gene: (a) CEU, GBR, FIN; (b) None; (c) FIN, CEU, GBR; JPT, CHB, CDX; (e) KHV, CDX, CHB, CHS, JPT; (f) KHV, JPT, CHS. The interval spanned by each gene is shaded in grey.

*SLC45A2* is a gene related to pigmentation (Branicki et al., 2008). It encodes a transporter protein that mediates melanin synthesis. In humans, it has been identified as a factor in the light skin of Europeans. As shown in Figure 6b, selection signals tended to be noisier in this region, and our median centered `btree` statistic does not see a pronounced peak this gene. The segments around this gene have small *p*-values for only TSI and CEU. However, the *p*-values are not above the genome-wide Bonferroni threshold, and are eclipsed by other nearby regions.

Figure 6c shows results for *HERC2*, which is associated with eye and skin pigmentation (Donnelly et al., 2012). Around this region there are blue-eye associated alleles found at high frequencies in European populations. In our results, the lowest *p*-value belongs to FIN, followed by GBR and CEU.

Turning to East Asian populations, we first studied *SLC44A5*, which is associated with neurological diseases and has been reported in several recent papers to be under selection in Japanese and Chinese populations (Liu et al., 2013; Zhao et al., 2019; Yasumizu et al., 2020). Our method confirms these findings (Figure 6), with highly significant hits centered on this gene for Japanese in Tokyo, Japan (JPT) and Han Chinese in Beijing, China (CHB).

We also found significant hits for all East Asian populations near *EDAR*(Figure 6e), again confirming earlier studies (Botchkarev and Fessing, 2005; Hlusko et al., 2018).

Finally, we examined the *ADH1* family. Alcohol is degraded primarily by alcohol dehydrogenase, and genetic variation affecting the rate of alcohol degradation found at *ADH1B* and *ADH1C*. Variants of these genes are thought to be associated with alcohol drinking habits and alcoholism. Our results (Figure 6f) confirm earlier findings (Han et al., 2007) that this family is under directional selection in Kinh in Ho Chi Minh City, Vietnam (KHV); Japanese in Tokyo, Japan (JPT); and Southern Han Chinese (CHS).

Estimates of the raw 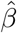 values corresponding to these Manhattan plots are given in the supplement (see Figures S6 and S7).

#### 3.3.2 Balancing selection

Next, we used our method to study long-term balancing selection in the the major histocompatibility complex (MHC). MHC is a large region of the vertebrate genome with immune-related functionality. Because evolution favors allelic diversity in this region (Takahata, 1993), we expect to detect signals of balancing selection in all populations. Our results (Figure 7) confirm this expectation; we observed highly significant signals across all 1000 Genomes subpopulations. Importantly, since this is an upper tail test for `bsfs`, we reject the null hypothesis that *β* = 0 in favor of the alternative *β* > 0. Thus, our method correctly infers that *MHC* is under balancing selection.

**Figure 7:**
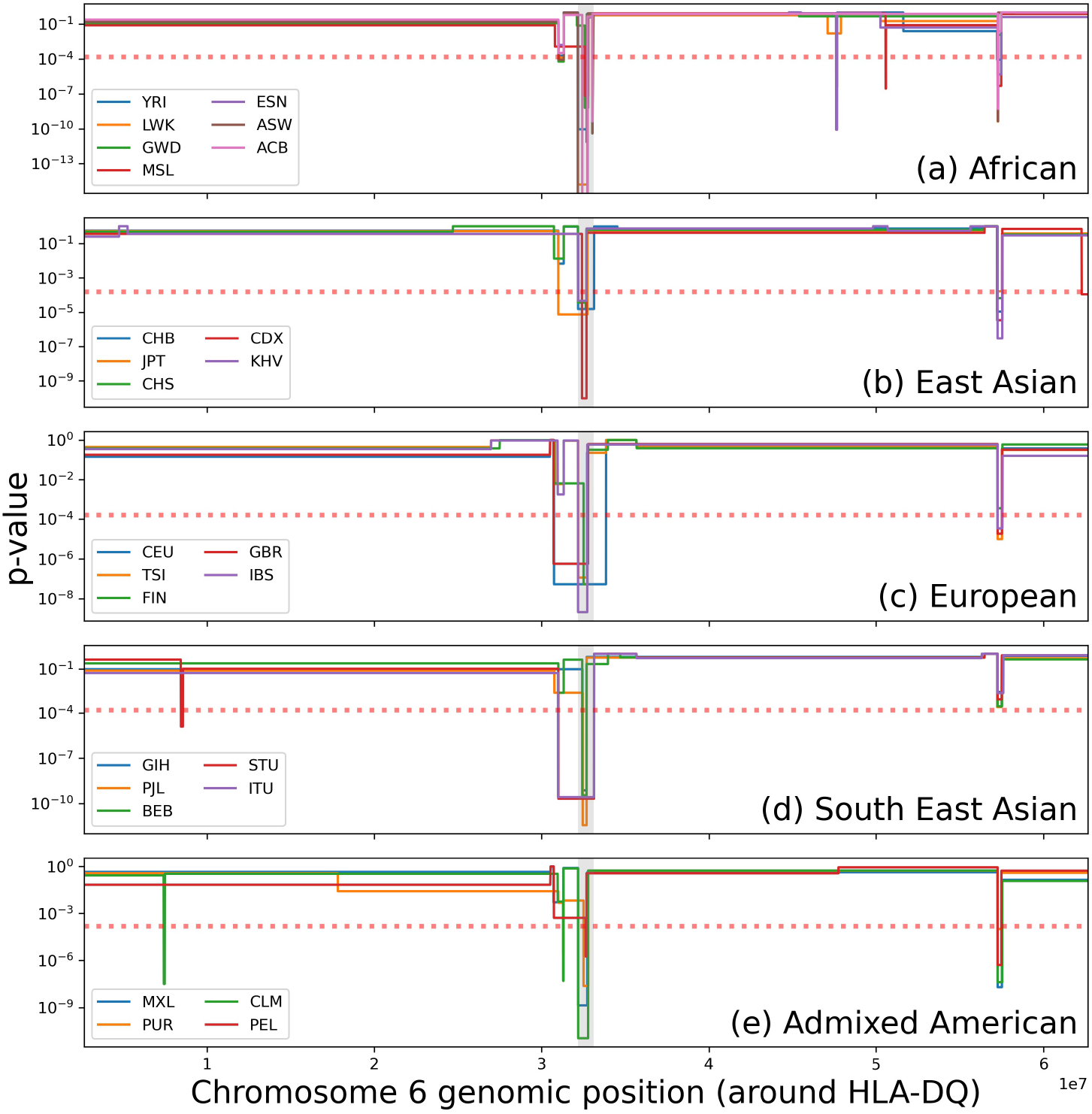
Genome Scan *p*-values of the bsfs segments around *HLA-DQ*. Most of the 1000 Genomes subpopulations have a pronounced balancing selection signal in this region.

#### 3.3.3 Results of genome-wide scan

In Section S5, we list the genomewide top hits (in terms of *p*-value) for the five major superpopulations in the 1000 Genomes dataset. They include a number of loci that are known to be under selection; such as *LCT*, *ALDH* ; the *HLA* complex; and various pigmentation, and eye color-related genes. There are also other hits that, as far as we can tell, have not yet been implicated by natural selection. Note that, due to linkage, many more genes are tagged than are likely under selection, but the genes should be proximal to a selected locus. A browser which can be used to explore all of our results, and compare them with classical tests of neutrality, is provided at the URL shown below.

## 4 Discussion

In this paper, we presented some new methods to detect natural selection using a generalization of Kingman’s coalescent to the case where genealogies exhibit systematic topological imbalance. We showed how this leads to relatively simple estimators of selection that can be applied to frequency spectrum data, or just as easily to sequences of estimated genealogies. An important feature of our method is its ability to incorporate demographic information. Using simulations, we recapitulated the tendency, already well known in the literature, of widely used deviance statistics like Tajima’s *D* to conflate variations in effective population size with natural selection. We showed that our method can correct for this tendency, by incorporating demographic estimates into its generative model of tree formation.

Our method is an example, albeit a basic one, of a recent trend towards likelihood-based methods for inferring natural selection from polymorphism data. We stress that our method will generally not be as sensitive as more elaborate and correct approximations to the coalescent under selection— compare, for example, the results of our Figures 2 and 3 with Figures 3 and 4 of Stern, Wilton, and Nielsen (2019). However, an advantage of our method is easy to understand and interpret, and also fast, requiring only to solve a univariate optimization problem. This can be done in only fractions of a second even for large sample sizes (Figure S3). Running our method on the entire 1000 Genomes dataset takes a few hours on a cluster. We see our work as adding to the toolbox of exploratory procedures that the analyst performs when studying a new dataset. Large “hits” yielded from our method can be used to flag a region for subsequent analysis, perhaps using more advanced and computationally expensive full-likelihood procedures. To this end, we have created an open source software package that makes it easy to run our methods. Researchers can also access our 1000 Genomes Project results with the browser we developed for this purpose. It enables to search through whole genome hits of our *β* estimates along with classical neutrality tests across for all populations.

There are several ways our model could be improved. We focused on the beta-binomial distribution because of its earlier usages in phylogenetics. However, as noted earlier, other distributions are possible, and perhaps some other model produces tree topology distributions that are more suited to studying natural selection. Another obvious criticism of our model is that it assumes that *β* is constant over time. This seems most appropriate for highly variable regions like *HLA*, where there is a continual introduction of new selected alleles. For regions that came under sudden directional selection as the result of the introduction of a beneficial allele, it would be better to use a model where the topological distribution of subtrees varies over time. This could allow for estimating the age of a selected variant, or understanding whether selection occurred on standing variation or because of the introduction of a new allele, both topics of longstanding interest in population genetics (Malaspinas et al., 2012; Hedrick, 2013; Barrett and Schluter, 2008; Feder, Kryazhimskiy, and Plotkin, 2014; Terhorst, Schlötterer, and Song, 2015; Palamara et al., 2018). Incorporating this feature into our SFS-based model would be challenging, as it creates dependence between the “time” and “topology” components of the expected frequency spectrum, thus invalidating equation (6). But it is easily added to the tree-based estimator in Section 2.3.2. We experimented with this, but found that the branch lengths from the current generation of tree sequence estimation programs are not yet reliable enough to support this kind of inference. As these methods continue to improve, this could be a future extension of our work.

On a similar note, when running our method on tree sequence data, we observed that the estimated trees contained many polytomies. Since trees generated under Kingman’s coalescent are almost surely bifurcating, we broke these polytomies arbitrarily in order to perform inference. However, polytomies could comprise another signal of selection, particularly in the case of recent positive selection. Incorporating a probabilistic model of node size into our method could potentially make use of this signal. The Λ-coalescent (Sagitov, 1999; Pitman, 1999) is a generalization of King-man’s coalescent which allows for various forms of multiple-merger events. Research on inference methods under generalized coalescents is ongoing (Spence, Kamm, and Song, 2016; Blath et al., 2016). In the future, our method could be extended to work under this more general model.

## Acknowledgements

This project was conceived during the workshop *Ecosystem dynamics: stakes, data and models* at Université Paris–Saclay. JT thanks the Institut Pascal for organizing the event, travel funding, and the limitless supply of M&M’s; and Amandine Véber and Raazesh Sainudiin for helpful early discussions. JT was supported by NSF grant DMS-2052653.

## Data and code availability

All of the data analyzed in this paper are publicly available. An open source implementation of our methods is available at https://github.com/jthlab/bim. Notebooks which reproduce our analyses are available at https://github.com/jthlab/bim-paper.

---

**Algorithm 1:**
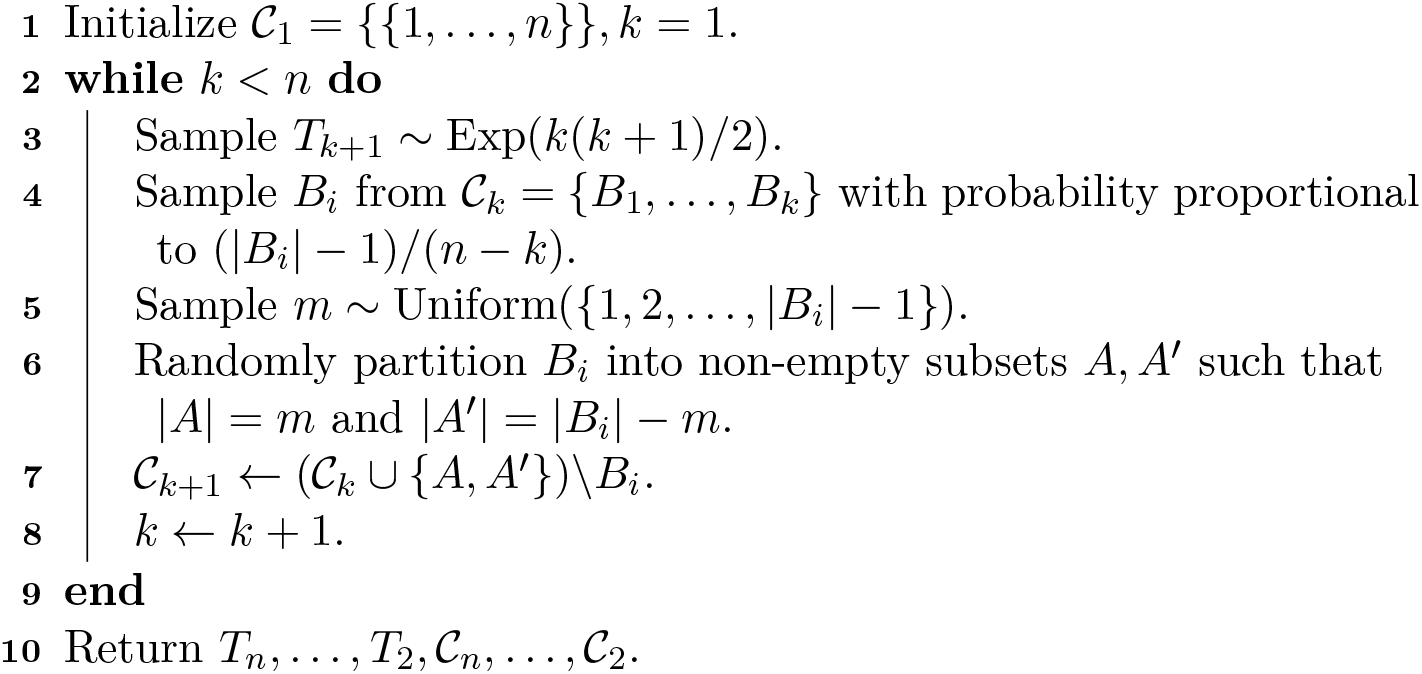
Kingman’s coalescent (forward-time version).

**Algorithm 2:**
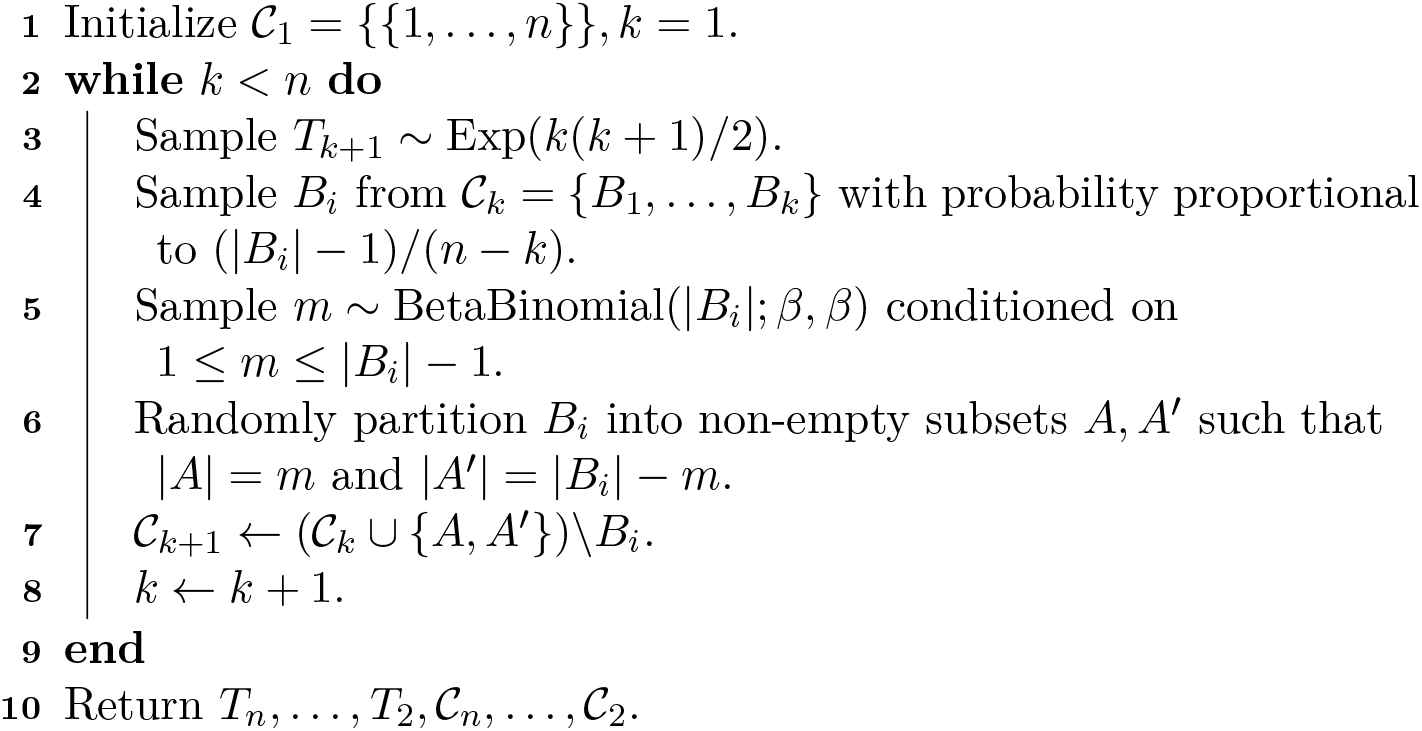
*β*-splitting coalescent model.

## Supplementary Materials

### S1 Data analysis pipeline

In the data analysis we will refer SFS-based *β*-splitting estimate as `bsfs` and tree-based *β*-splitting estimate as `btree`.

#### S1.1 Simulation studies

All simulations were performed using SLiMv3 (Haller and Messer, 2019). An example for the complete simulation pipeline can be seen in Figure S1.

(a) For each simulation, we have a set of parameters *θ*, and we run that set 250 iterations with a different seed.
(b) In SLiM we drop non-neutral mutations and run it until a stopping condition, and then record the tree-sequence.
(c) SLiM is a forward simulator and does not guarantee a common ancestor. After the simulation is completed, genealogies with multiple roots are recapitated using the procedure described by Haller et al. (2019), and neutral mutations are introduced.
(d) bsfs is inferred from the allele frequency spectrum. For btree we estimated tree-sequences from the genotype matrix using using tsinfer (Kelleher et al., 2019a).
(e) A simulated region covers many trees (because of the recombination).

In order to represent the region, we combine these `btree`’s by taking span weighted averages. An example calculation is shown in Figure S1e. There are four `btree` estimates in this 10kb region, each spanning a different non-overlapping region. For this simulation `btree` value is calculated as:

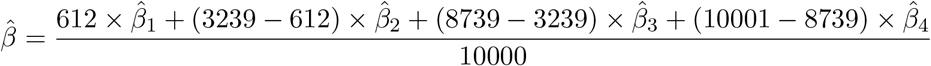

#### S1.2 Real data analysis

We tested our model on 1000 Genomes Project. A toy example of our analysis pipeline can be seen in Figure S2. For each 26 populations we repeat the following process:

(a) The genome-wide SFS is computed for each subpopulation.
(b) Population-specific size histories *η* are inferred using these empirical frequency spectra.
(c) For SFS-based analysis, genomes are divided into intervals as described below, and the local frequency spectrum is calculated for each interval conditional on the size history function inferred in the previous step.
(d) Tree-sequences subdivide the genome into disjoint regions spanned by each local tree.
(e) SFS-based estimates are obtained by estimating bsfs using a sliding window along the genome.
(f) Tree-sequence estimates are performed for each local genealogy.
(g) Tree-based estimates are converted to genomic coordinates using the averaging procedure described above.
(h) To combine these spatially correlated estimates, we use a changepoint detection procedure to aggregate the `bsfs` and `btree` estimates into a set of piecewise-constant functions (Celisse et al., 2018; Truong, Oudre, and Vayatis, 2020). Heuristically, we chose a fixed number of change-points for all populations. We chose in a way that we expect a change point on the average of 10Mb (3,100Mb/10Mb = 310 change points).
(i) Finally, *p*-values are computed for each segment using the procedure described in the Section S1.2.3.

Genomic scan statistics depend on the choice of window size, and stride (number of base pairs between the start of each consecutive window). Since we are using estimated tree-sequences for 26 populations, the start and end points of these estimated trees are different in each population. In real data analysis we constructed two different methods of window statistics for directional selection and balancing selection. For directional selection, we are seeking population specific signal of the selection (that is the reason why we use a median-centered *β* estimate, Section S1.2.1). But in order to compare `btree` statistic with `bsfs` (and other neutrality tests) within the population and with other `btree`’s across the populations we need them to be defined on the same positions on the genome. To overcome this we first estimate tree-based statistics (`btree` and Colless) for each tree. Second we define the windows in base pairs (we defined window size 10kbp and stride size 5kbp) and calculate SFS-based statistics (`bsfs` and other neutrality tests) for these windows. Finally we take averages of tree-based statistic inside each windows (See Section S1.1 part (e)). In order to detect long term balancing selection, we do not need the windows to be defined in the same regions. Since we are not looking for a population specific signal. After some trial and error, we see that windows defined by estimated tree locations give better results for balancing selection. Instead of sliding base pairs, we chose window size of 64 trees, and a stride of 32 trees, we calculated the SFS-based statistics on those regions.

In directional selection setup, for `bsfs`, this window-size and stride setup resulted 520,940 windows along the human genome for `bsfs`. This required us to solve 520940×26 = 13, 544, 440 optimization problems. For `btree`, each population has different number of trees, and in total it resulted 142,637,760 optimization tasks. Together with other statistics, all calculations required less than four hours on a cluster. For the balancing selection setup, number of tree estimates doesn’t change so we do not need to estimate `btree` again. But we estimated `bsfs` again for the different windows which required solving an additional 8, 914, 860 optimization problems and this finished approximately in 2 hours.

##### S1.2.1 Median-centered estimates of *β*

Some regions on the chromosome experience the similar evolutionary history among all human populations. For some regions, this causes a pronounced spike in |*β*| for all 26 populations in 1000 Genomes Project data, confounding our ability to detect population-specific signals of selection. To correct for this, we performed median-centering for windowed statistic: let 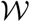 defines the set of windows along the chromosome, and 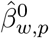 be the *β* estimate of window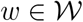 for each of the 26 1kg subpopulations

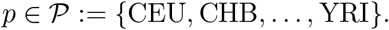

Then the median-centered estimate of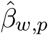 is defined as

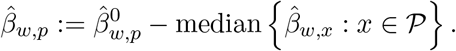

##### S1.2.2 Combining multiple 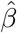 in each segments

Each segment decided by change point detection spans multiple windows. We average these windowed estimates of *β* and get a single estimate that represents each segment. To combine window estimates we used weighted averaging. Given *n_g_* windows spanning a given segment, we define 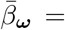 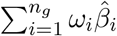 to be a weighted sum of estimated 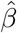 parameters, where 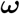is a weight vector. After some experimenting, we found that choosing the entries of 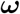 to be proportional to the proximity of the mid-point of the segment worked well, and all reported results are based on that choice of weights.

##### S1.2.3 Significance testing

Since the 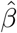are maximum likelihood estimates, 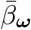 has approximately a normal distribution. To form *p*-values we therefore require the mean and variance of this statistic under the null hypothesis. Under neutrality, 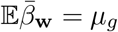, where *µ_g_* ≈ 0 is the chromosome-wide average which is determined empirically. To calculate 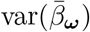 we need to consider the dependence between sequential estimates, which is non-zero due to linkage. We define

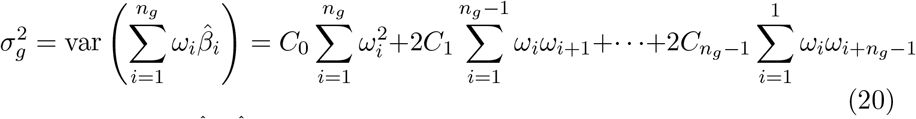

where 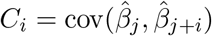 is the lag-*i* autocovariance term, which is assumed to be stationary (does not depend on *j*). The coefficients *C_i_* were are esti-mated empirically using chromosome-wide averages.

For each location we perform a one-sided test to determine whether 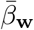 is abnormally high (signifying balancing selection) or low (directional selection). The corresponding *p*-values are

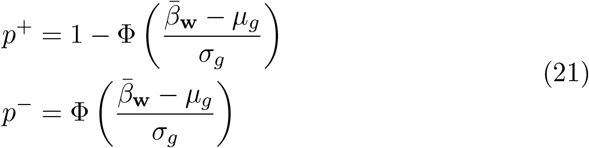

Finally, the *p*-values are Bonferroni corrected to account for multiple testing. In this case for 1000 genomes project population, we defined 311 segments, then the significance level will be equal to 0.05/311 ≈ 1.6 × 10^−4^.

## S2 Supplemental algorithms

**Algorithm S1:**
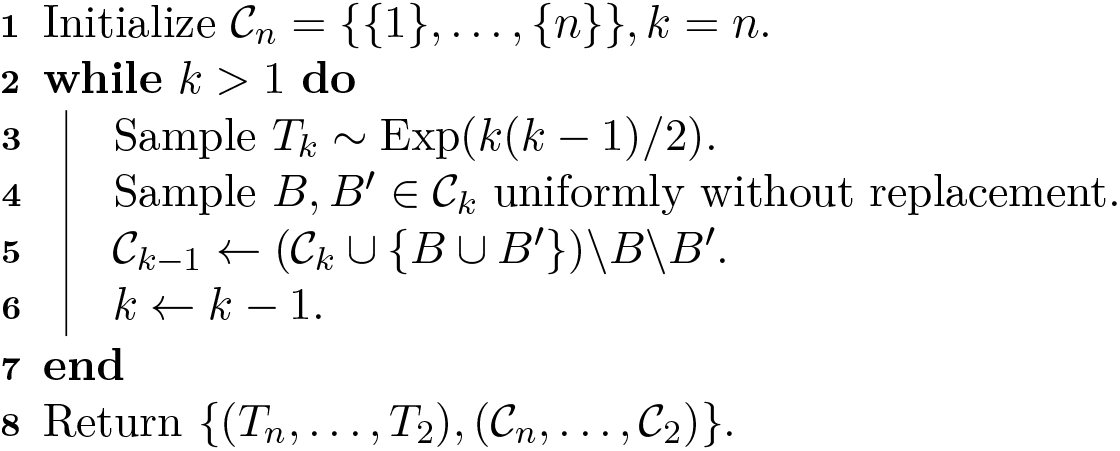
Kingman’s coalescent.

## S3 Simulation settings and code

**Table S1:** Simulation settings for directional selection experiment. *N_e_*(0) is population size at the start of the simulation, *g* is the growth rate of exponential growth, *t_g_* is the generations ago when exponential growth starts prior to sampling, *s* is the selective coefficient of the beneficial mutation and *h* is the dominance coefficient.

| Scenario | $N_e(0)$ | $g$ | $t_g$ | $s$ | $h$ |
| --- | --- | --- | --- | --- | --- |
| 1 | $2 \times 10^3$ | 0 | – | 0 | 0.5 |
| 2 | $10^3$ | 0.01 | 250 | 0 | 0.5 |
| 3 | $2 \times 10^3$ | 0 | – | 0.05 | 0.5 |
| 4 | $10^3$ | 0.01 | 250 | 0.05 | 0.5 |

**Table S2:** Simulation settings for balancing selection experiment. *µ_s_* is the mutation rate for advantageous mutation, *t_m_* is the number of generations age that selection began, and the rest of the parameters are as in Table S1.

| Scenario | $N_e(0)$ | $\mu_s$ | $t_m$ | $g$ | $t_g$ | $s$ | $h$ |
| --- | --- | --- | --- | --- | --- | --- | --- |
| 1 | $2 \times 10^3$ | 0 | – | 0 | – | 0 | 0.5 |
| 2 | $10^3$ | 0 | – | 0.01 | 250 | 0 | 0.5 |
| 3 | $2 \times 10^3$ | $10^{-8}$ | 4250 | 0 | – | 0.002 | 25 |
| 4 | $10^3$ | $10^{-8}$ | 4250 | 0.01 | 250 | 0.002 | 25 |

**S3.1.**
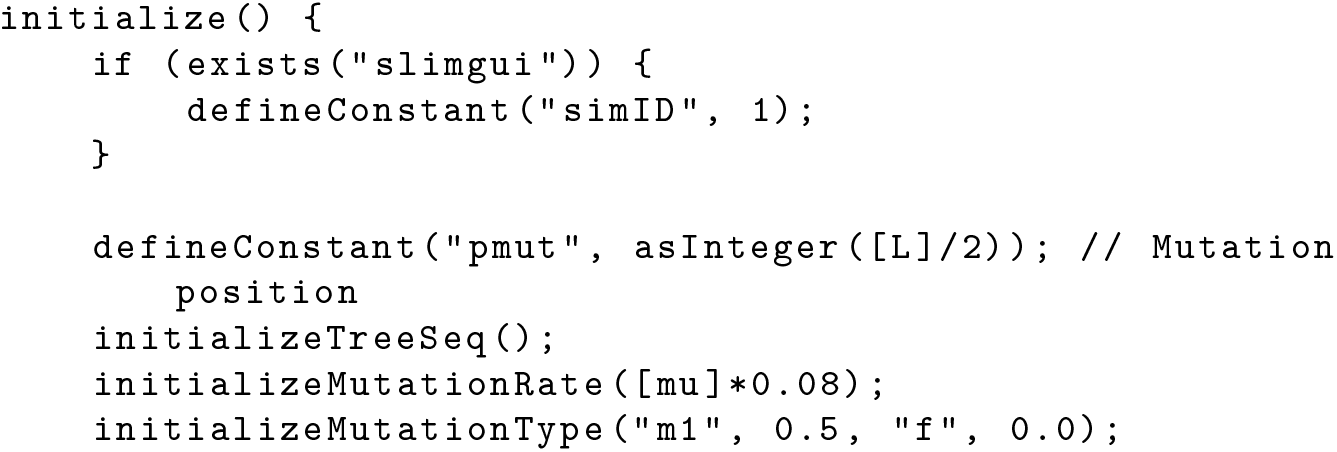
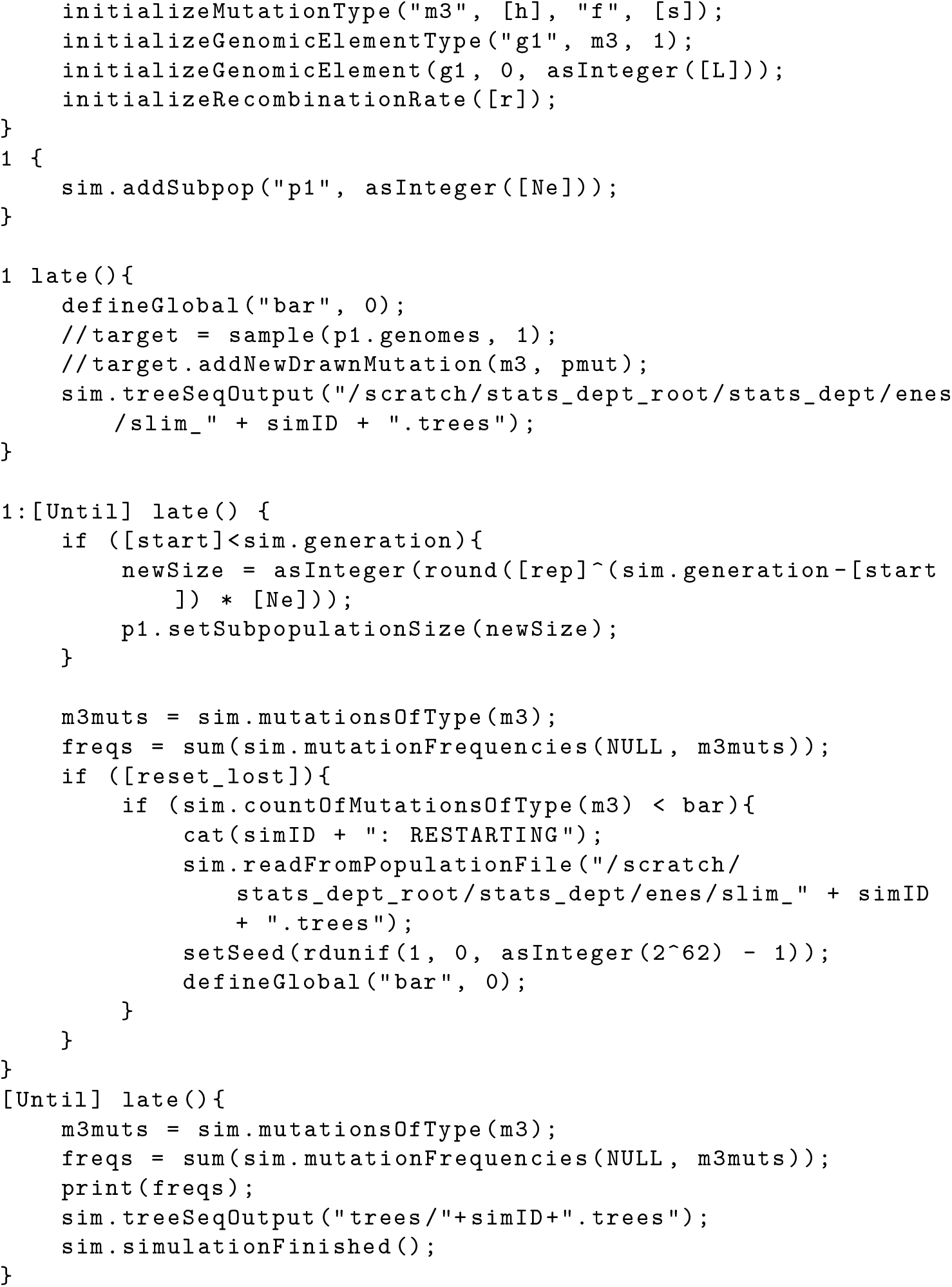
Directional selection.

**S3.2.**
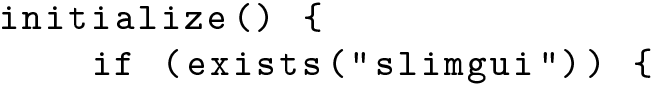
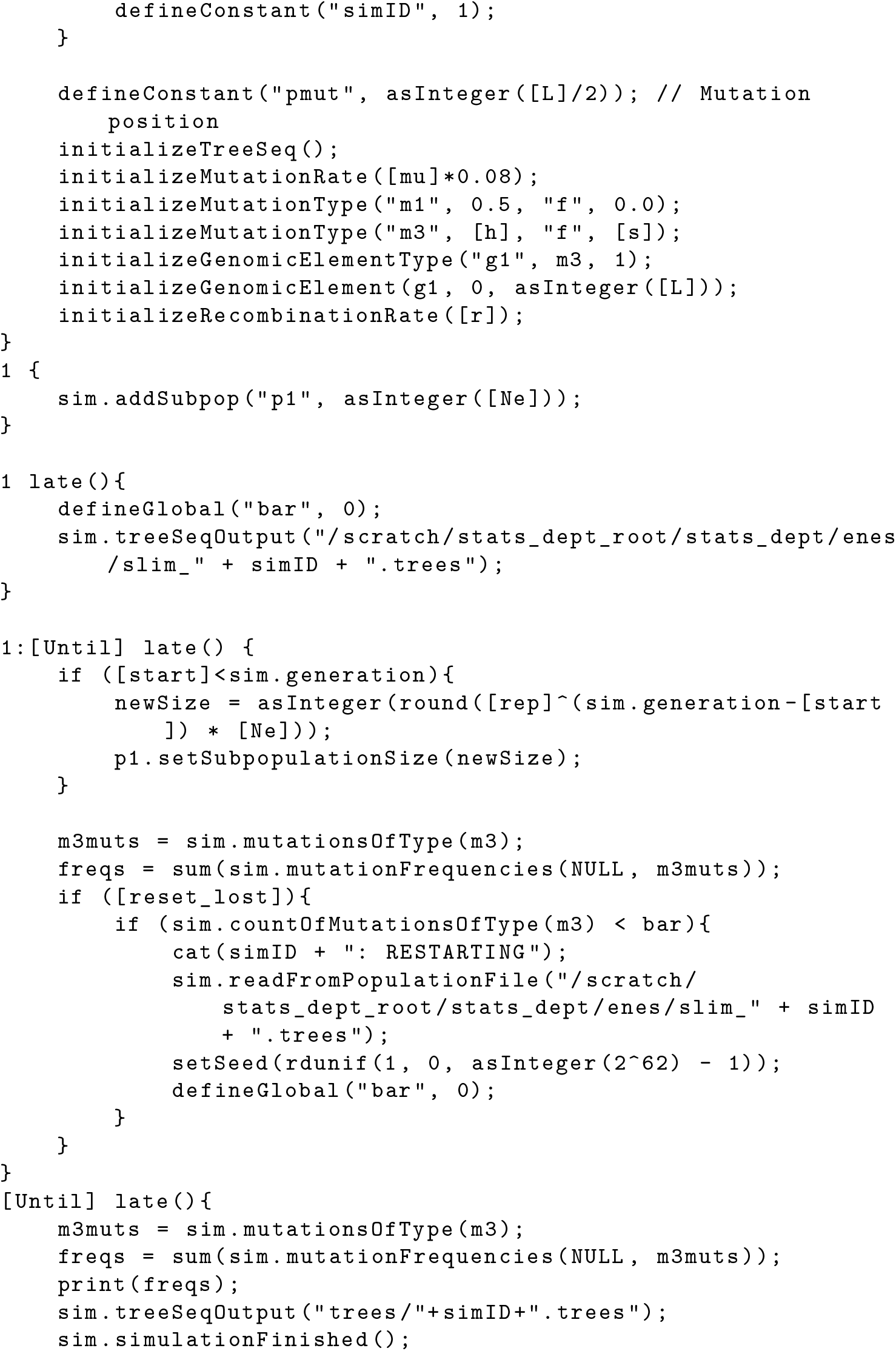
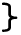
Balancing selection.

## S4 Supplemental figures

**Figure S1:**
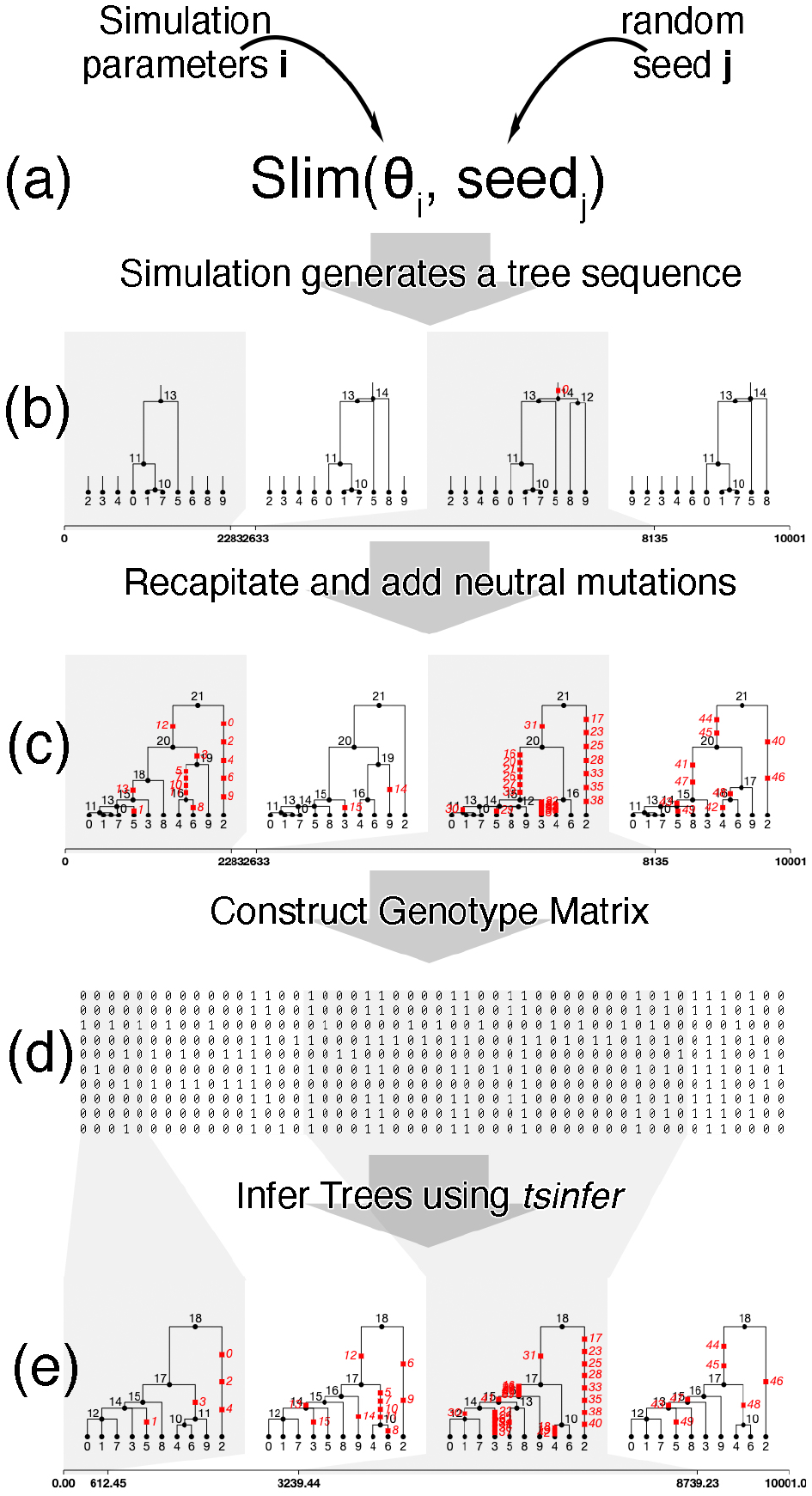
Simulation pipeline. (a) In each set of simulations we fix a group of parameters (*θ*) and a random seed. (b) After simulation is finished, we save the tree sequence of the true genealogy. It can be seen that trees have multiple roots and the sample only have one mutation at the third tree. (c) We recapitate the trees and add the neutral mutations. (d) Genotype matrix is extracted from the tree sequence and used to estimate `bsfs`. (e) Tree sequences are inferred from genotype matrices and used to estimate btree.

**Figure S2:**
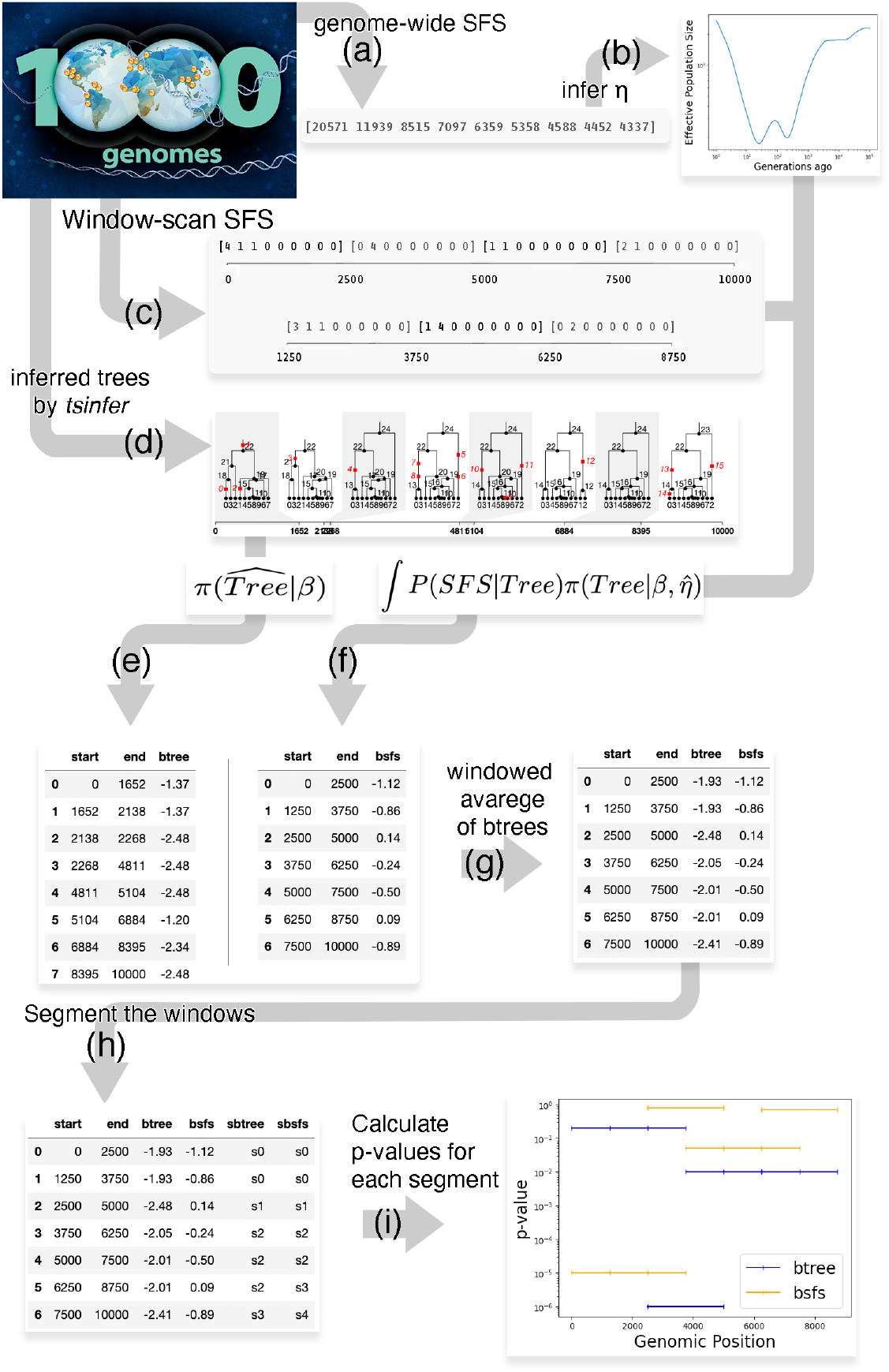
Real data analysis pipeline. (a) We calculate genome-wide SFS for the sample. (b) Using this we infer population size histories (*η*). (c) We calculate windowed statistic of SFS. Window size is 2500 and stride is 1250. With the region of size 10000, we get 7 windows. (d) From genotype matrix, we infer trees using `tsinfer`. (d) For each tree in the tree sequence, we estimate `btree`. (f) For each windowed-statistic SFS we estimate `bsfs`. (g) We take average `btree`s inside bsfs regions by taking span weighted avarage of each. (h) We apply the change point detection method to define segments. (i) We calculate the p-values for each segment.

**Figure S3:**
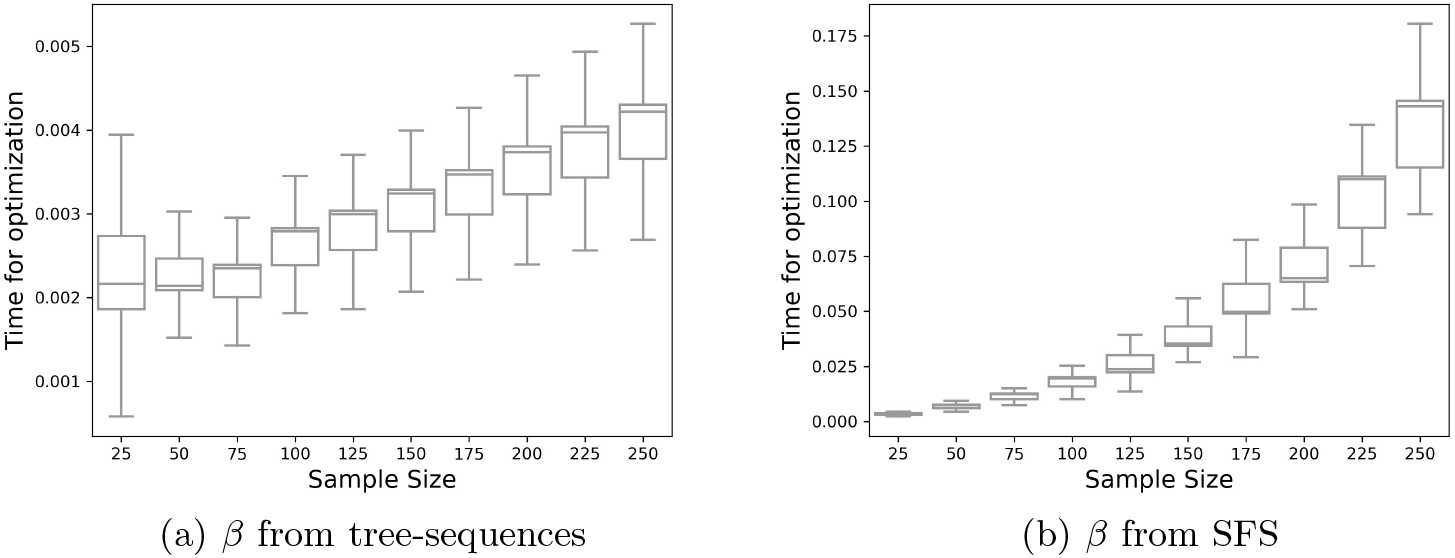
Time-complexity of estimating *β* for a single optimization task.

**Figure S4:**
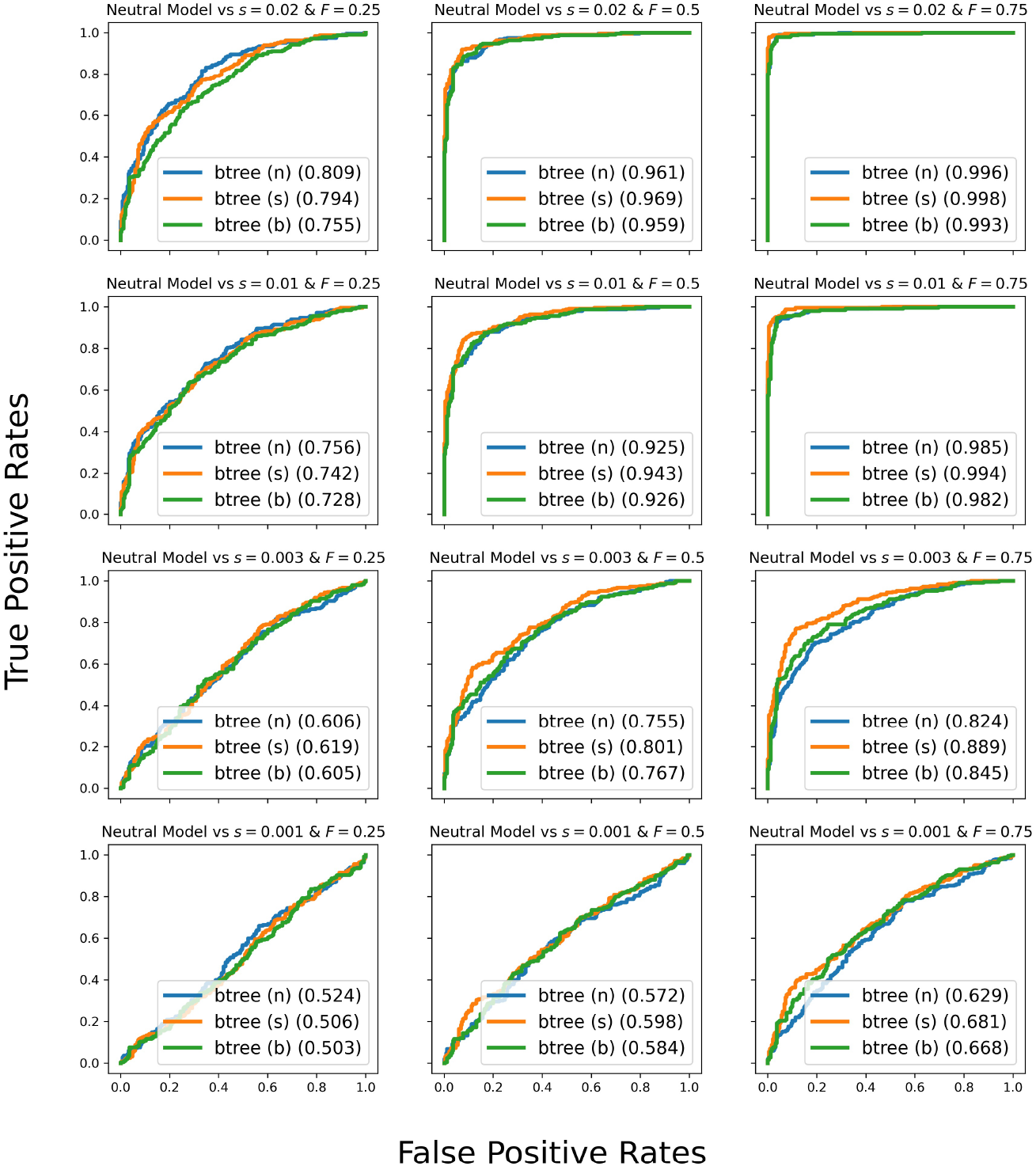
ROC curves for positive genic selection using `btree` with three different choices of likelihood weights. *s* represents selective advantage of the mutation and *F* represents allele frequency of the mutation in the sample. Letter inside the parenthesis represents the likelihood weighting method; (n):no weighting, (s): size of the internal node, (b): total amount of branch length

**Figure S5:**
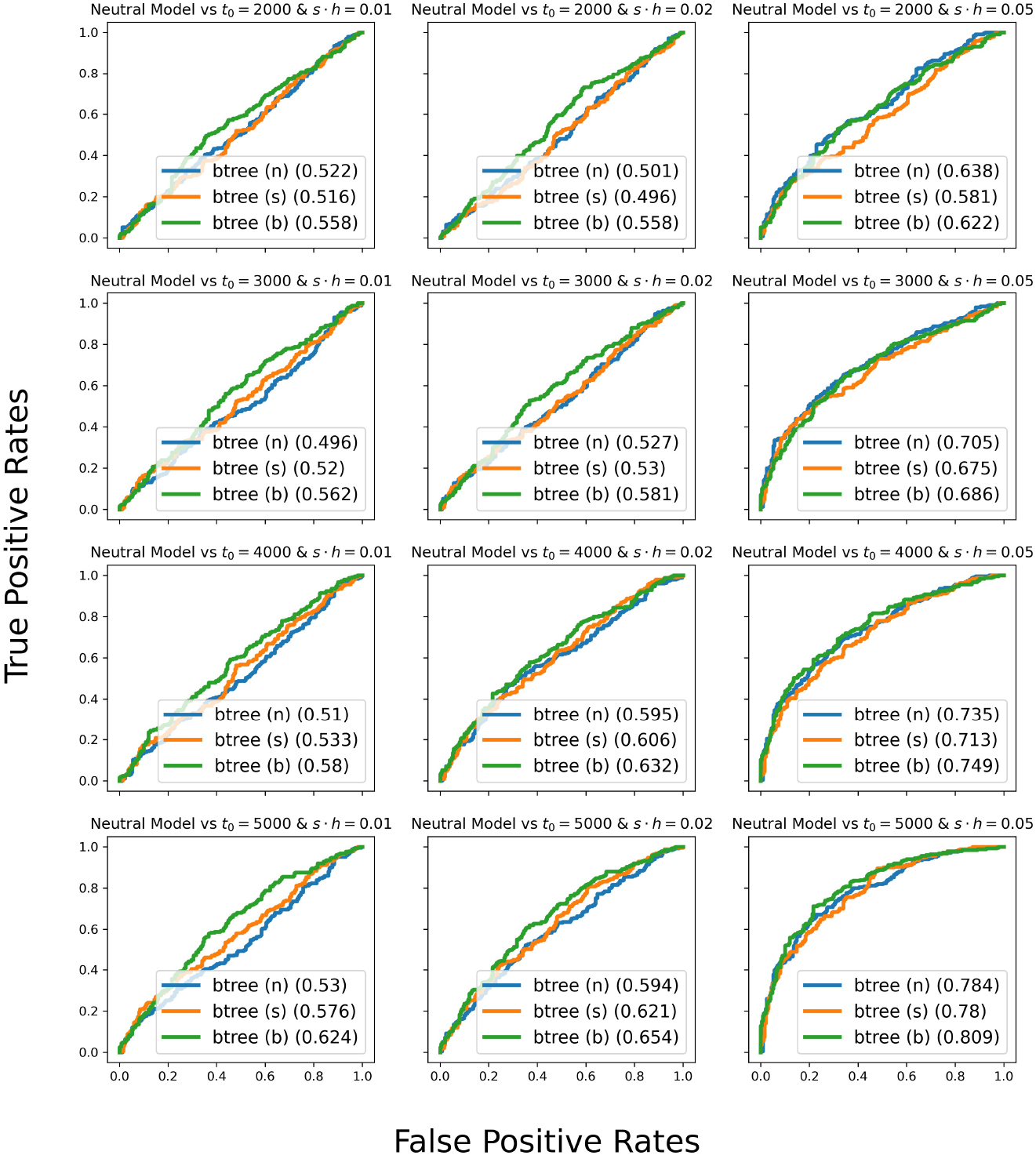
ROC curves for advantageous heterozygote mutation simulations using `btree` with three different choices of likelihood weights. *s* represents selective advantage of the mutation, *h* is the dominance factor. *t*_0_ represents how many generations ago the advantageous mutations were introduced into the sample. Letter inside the parenthesis represents the likelihood weighting method; (n):no weighting, (s): size of the internal node, (b): total amount of branch length

**Figure S6:**
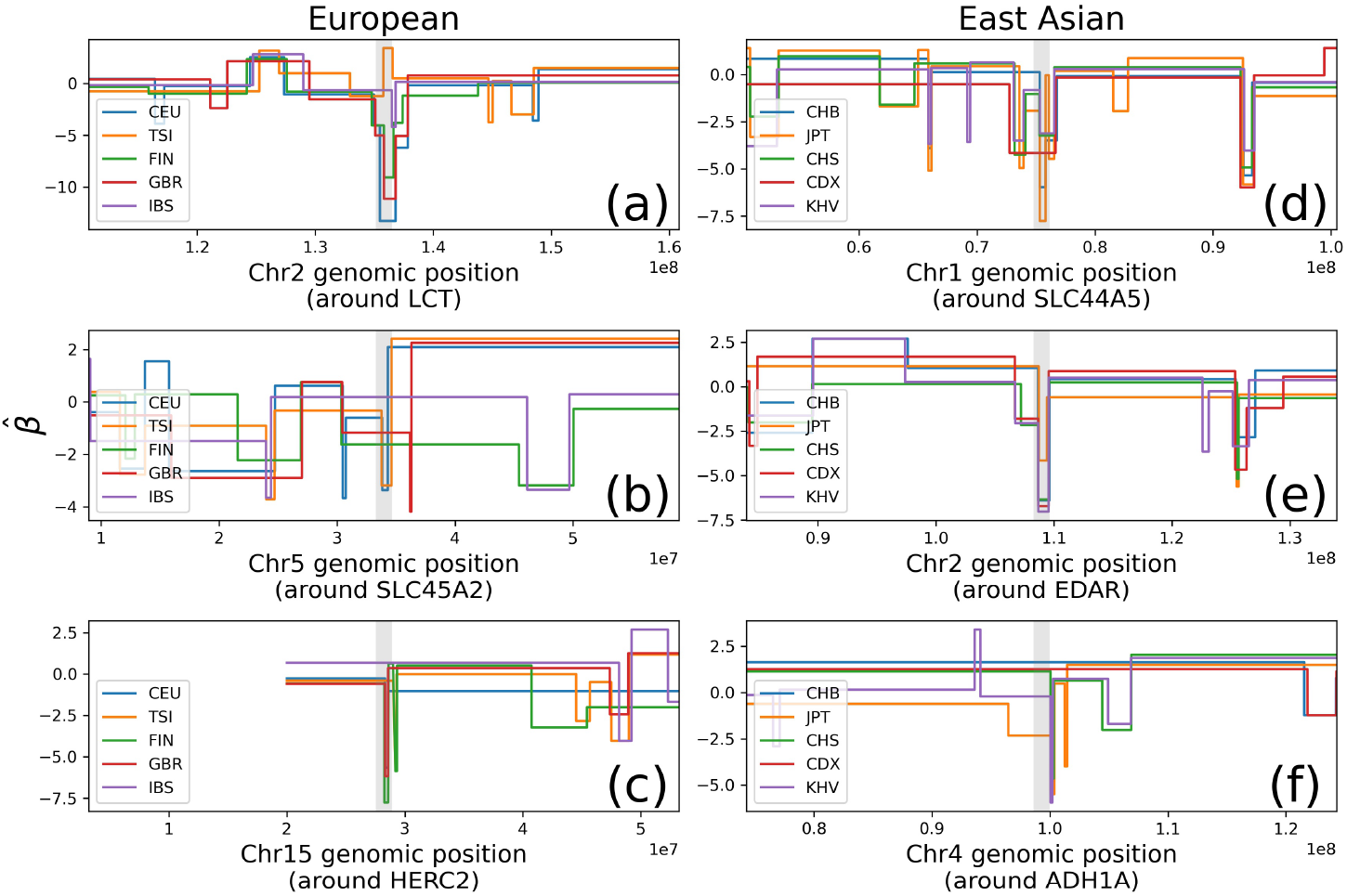
Directional selection examples for 1000 Genomes Project. Segmented genome-scan *β* estimates by median centered `btree` are provided. See Figure 6.

**Figure S7:**
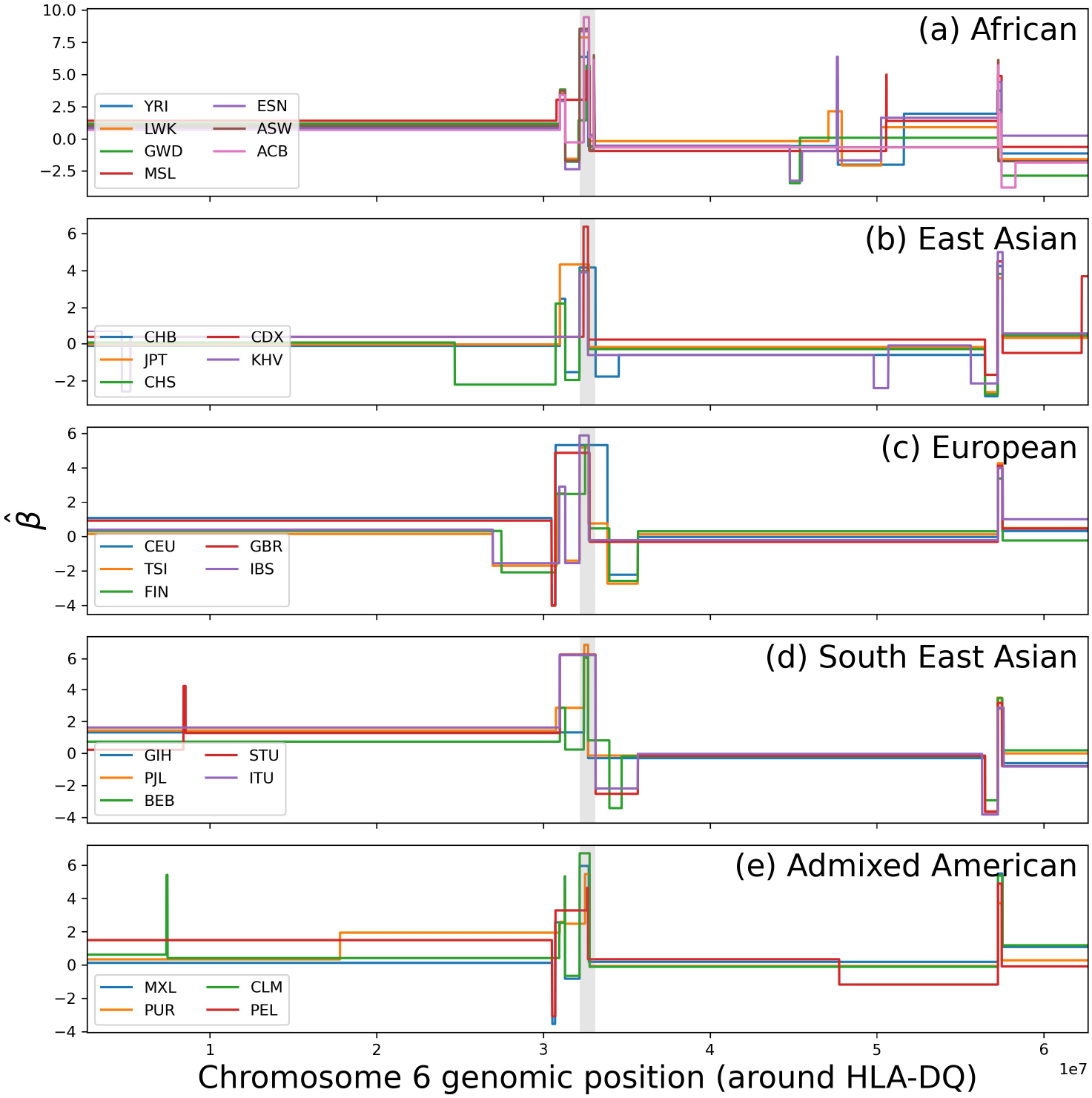
Genome Scan *β* estimates by `bsfs` segments around *HLA-DQ*. See Figure 7.

## S5 Supplemental Tables

**Table S3:**
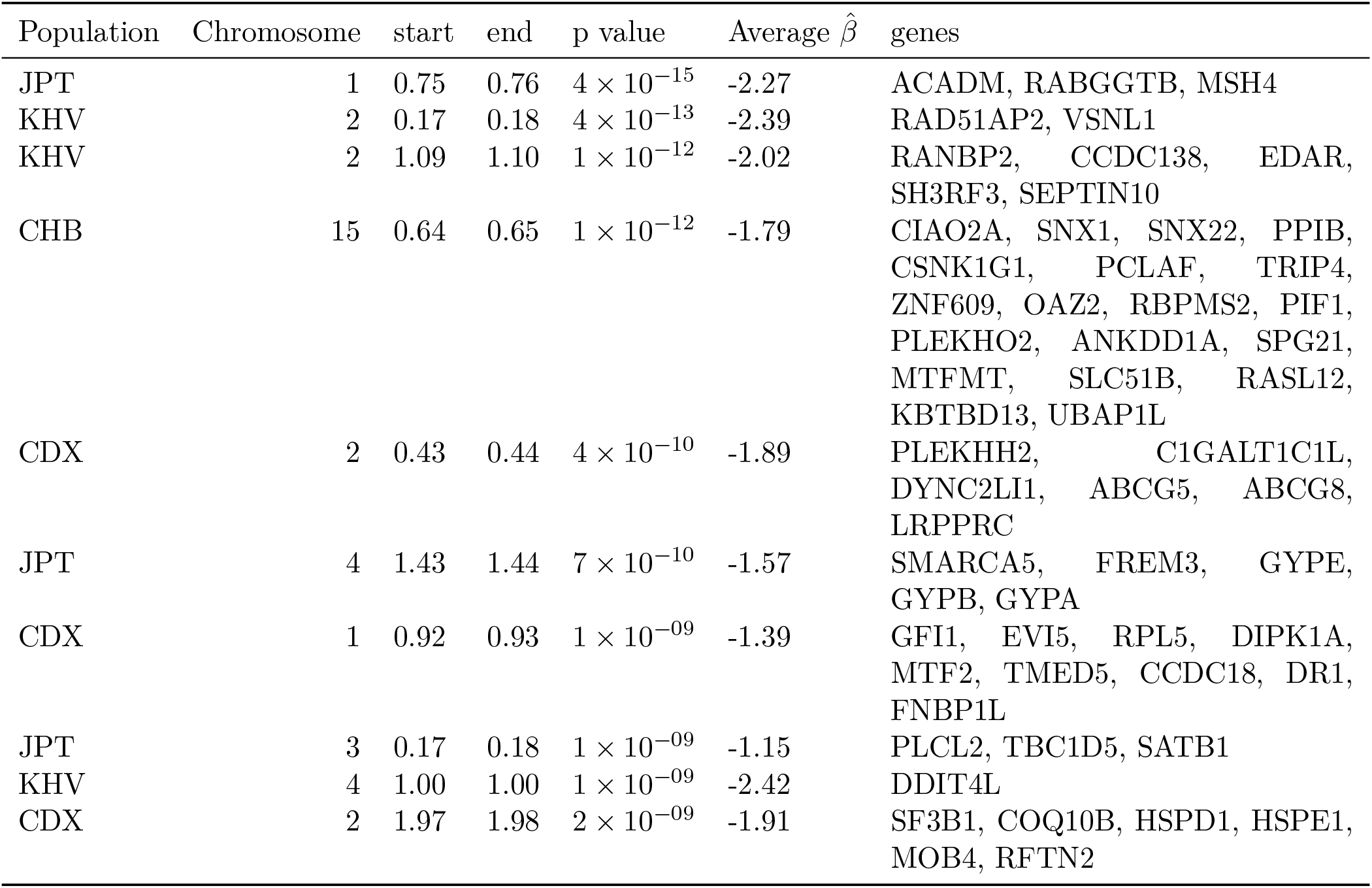
Most significant directional selection hits for East Asian

**Table S4:** Most significant directional selection hits for European

| Population | Chromosome | start | end | p value | Average $\hat{\beta}$ | genes |
| --- | --- | --- | --- | --- | --- | --- |
| CEU | 2 | 1.35 | 1.37 | $2 \times 10^{-40}$ | -2.31 | R3HDM1, UBXN4, LCT, MCM6, DARS1, CXCR4, THSD7B |
| GBR | 12 | 1.11 | 1.13 | $3 \times 10^{-18}$ | -1.50 | ATXN2, BRAP, ACAD10, ALDH2, MAPKAPK5, TMEM116, ERP29, NAA25, TRAFD1, RPL6, PTPN11, RPH3A |
| FIN | 15 | 0.28 | 0.29 | $4 \times 10^{-15}$ | -1.73 | GOLGA8F, GOLGA8G |
| TSI | 11 | 0.89 | 0.89 | $5 \times 10^{-13}$ | -1.63 | TYR |
| TSI | 5 | 1.30 | 1.31 | $4 \times 10^{-11}$ | -1.33 | CHSY3, HINT1, LYRM7, CDC42SE2 |
| GBR | 8 | 0.12 | 0.13 | $5 \times 10^{-11}$ | -1.98 | DEFB130A, FAM86B2 |
| FIN | 6 | 0.28 | 0.29 | $9 \times 10^{-11}$ | -1.35 | GPX5, ZBED9 |
| GBR | 15 | 0.75 | 0.75 | $3 \times 10^{-10}$ | -1.39 | CLK3, EDC3, CYP1A1, CYP1A2, CSK, LMAN1L, CPLX3, ULK3, SCAMP2, MPI, FAM219B, COX5A, RPP25, SCAMP5, PPCDC |
| GBR | 6 | 0.35 | 0.35 | $8 \times 10^{-10}$ | -1.42 | TCP11, SCUBE3, ZNF76, DEF6, PPARD |
| CEU | 1 | 1.00 | 1.00 | $1 \times 10^{-09}$ | -1.87 | AGL |

**Table S5:** Most significant directional selection hits for African

| Population | Chromosome | start | end | p value | Average $\hat{\beta}$ | genes |
| --- | --- | --- | --- | --- | --- | --- |
| ESN | 19 | 0.43 | 0.44 | $4 \times 10^{-17}$ | -2.19 | PSG5, PSG4, PSG9, CD177, TEX101, LYPD3, PHLDB3, ETHE1, ZNF575, XRCC1 |
| ESN | 6 | 0.53 | 0.53 | $5 \times 10^{-16}$ | -2.74 | TMEM14A |
| GWD | 6 | 0.59 | 0.62 | $9 \times 10^{-14}$ | -2.25 | KHDRBS2 |
| LWK | 8 | 0.44 | 0.47 | $1 \times 10^{-12}$ | -1.60 | SPIDR |
| ESN | 11 | 0.49 | 0.49 | $2 \times 10^{-11}$ | -1.60 | TRIM49B, TRIM64C |
| YRI | 1 | 1.62 | 1.62 | $2 \times 10^{-11}$ | -1.80 | DUSP12, ATF6 |
| ESN | 1 | 1.54 | 1.54 | $7 \times 10^{-11}$ | -1.73 | CRTC2, SLC39A1, CREB3L4, JTB, RAB13, RPS27, NUP210L |
| ESN | 3 | 1.25 | 1.26 | $2 \times 10^{-9}$ | -1.61 | SNX4, OSBPL11 |
| ASW | 13 | 0.52 | 0.52 | $4 \times 10^{-9}$ | -1.43 | NEK3 |
| YRI | 19 | 0.39 | 0.39 | $4 \times 10^{-9}$ | -1.68 | LGALS7, LGALS7B, LGALS4, ECH1, HNRNPL, RINL, SIRT2 |

**Table S6:**
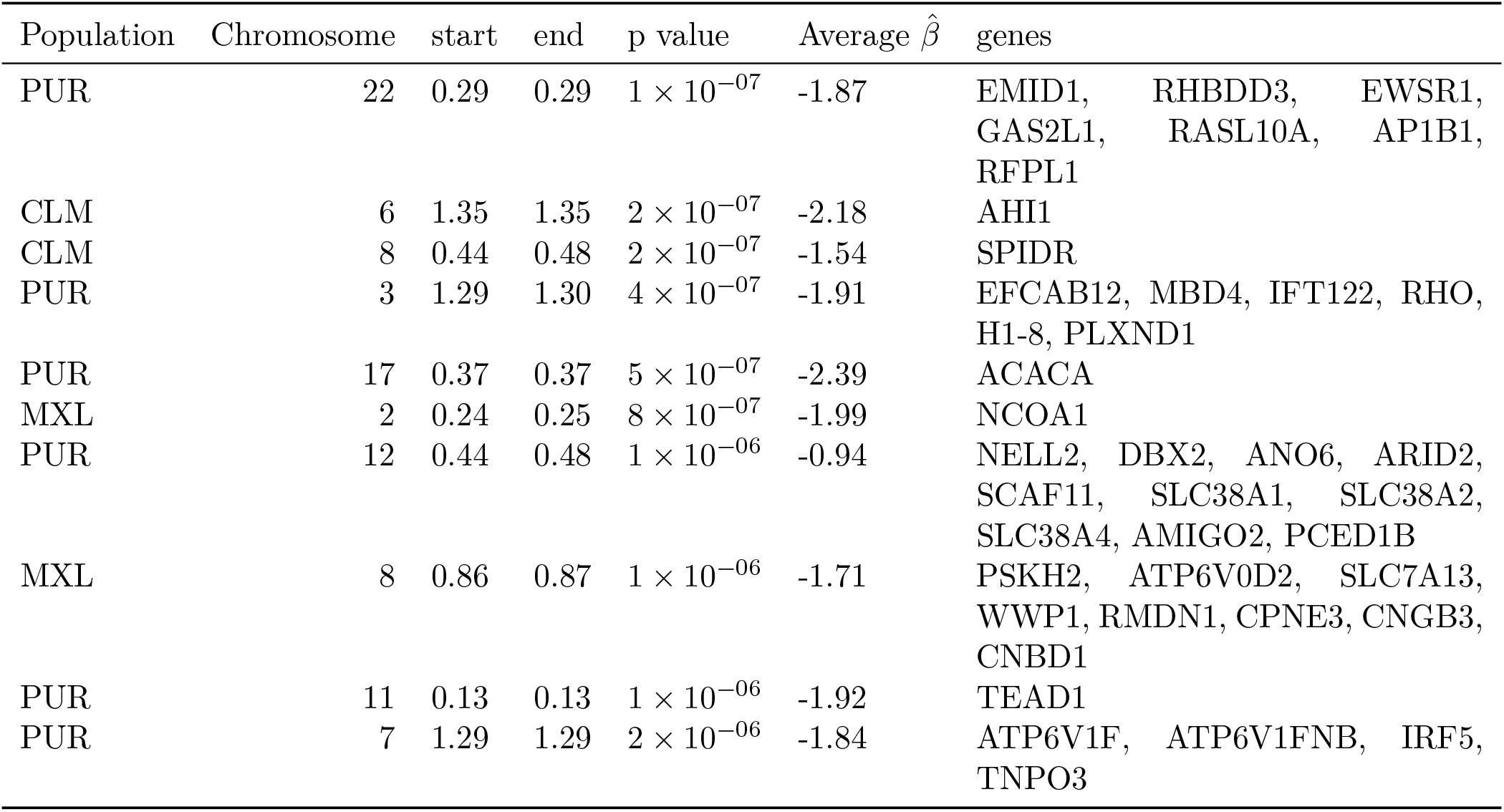
Most significant directional selection hits for Ad-Mixed American

| Population | Chromosome | start | end | p value | Average $\hat{\beta}$ | genes |
| --- | --- | --- | --- | --- | --- | --- |
| PUR | 22 | 0.29 | 0.29 | $1 \times 10^{-07}$ | -1.87 | EMID1, RHBDD3, EWSR1,<br>GAS2L1, RASL10A, AP1B1,<br>RFPL1 |
| CLM | 6 | 1.35 | 1.35 | $2 \times 10^{-07}$ | -2.18 | AHI1 |
| CLM | 8 | 0.44 | 0.48 | $2 \times 10^{-07}$ | -1.54 | SPIDR |
| PUR | 3 | 1.29 | 1.30 | $4 \times 10^{-07}$ | -1.91 | EFCAB12, MBD4, IFT122, RHO,<br>H1-8, PLXND1 |
| PUR | 17 | 0.37 | 0.37 | $5 \times 10^{-07}$ | -2.39 | ACACA |
| MXL | 2 | 0.24 | 0.25 | $8 \times 10^{-07}$ | -1.99 | NCOA1 |
| PUR | 12 | 0.44 | 0.48 | $1 \times 10^{-06}$ | -0.94 | NELL2, DBX2, ANO6, ARID2,<br>SCAF11, SLC38A1, SLC38A2,<br>SLC38A4, AMIGO2, PCED1B |
| MXL | 8 | 0.86 | 0.87 | $1 \times 10^{-06}$ | -1.71 | PSKH2, ATP6V0D2, SLC7A13,<br>WWP1, RMDN1, CPNE3, CNGB3,<br>CNBD1 |
| PUR | 11 | 0.13 | 0.13 | $1 \times 10^{-06}$ | -1.92 | TEAD1 |
| PUR | 7 | 1.29 | 1.29 | $2 \times 10^{-06}$ | -1.84 | ATP6V1F, ATP6V1FNB, IRF5,<br>TNPO3 |

**Table S7:** Most significant directional selection hits for South East Asian

| Population | Chromosome | start | end | p value | Average $\hat{\beta}$ | genes |
| --- | --- | --- | --- | --- | --- | --- |
| BEB | 16 | 0.34 | 0.47 | $3 \times 10^{-17}$ | -1.45 | SHCBP1, VPS35, ORC6, MYLK3, C16orf87, GPT2 |
| GIH | 15 | 0.48 | 0.49 | $3 \times 10^{-12}$ | -1.69 | DUT, FBN1 |
| STU | 17 | 0.44 | 0.44 | $3 \times 10^{-12}$ | -1.77 | HROB, ASB16, TMUB2, ATXN7L3, UBTF, SLC4A1, RUNDC3A, SLC25A39, GRN, FAM171A2 |
| GIH | 3 | 0.89 | 0.94 | $8 \times 10^{-12}$ | -0.84 | EPHA3, PROS1 |
| GIH | 2 | 0.97 | 0.97 | $1 \times 10^{-11}$ | -1.73 | FAHD2B |
| STU | 16 | 0.30 | 0.31 | $4 \times 10^{-11}$ | -1.23 | SEPHS2, ITGAL, ZNF768, ZNF747, ZNF764, ZNF688, ZNF785, ZNF689, PRR14, FBRS, SRCAP, TMEM265, PHKG2, CCDC189, RNF40, ZNF629, BCL7C, CTF1, FBXL19, ORAI3, SETD1A, HSD3B7, STX1B, STX4, ZNF668, ZNF646, PRSS53, VKORC1, BCKDK, KAT8, PRSS8, PRSS36, FUS, PYCARD, TRIM72, PYDC1, ITGAM, ITGAX, ITGAD, COX6A2, ZNF843, ARMC5, TGFB1I1 |
| PJL | 2 | 2.23 | 2.23 | $5 \times 10^{-11}$ | -1.60 | FARSB |
| PJL | 2 | 0.92 | 0.95 | $6 \times 10^{-11}$ | -1.63 | TEKT4, MAL, MRPS5, ZNF514, ZNF2, PROM2, KCNIP3 |
| STU | 10 | 0.93 | 0.94 | $1 \times 10^{-10}$ | -1.20 | CEP55, FFAR4, RBP4, PDE6C, FRA10AC1, LGI1, SLC35G1, PLCE1 |
| STU | 19 | 0.36 | 0.36 | $4 \times 10^{-10}$ | -1.90 | ZFP82 |

**Table S8:** Most significant directional selection hits for 1000 Genomes Project Populations

| Population | Chromosome | start | end | p value | Average $\hat{\beta}$ | genes |
| --- | --- | --- | --- | --- | --- | --- |
| CEU | 2 | 1.35 | 1.37 | $2 \times 10^{-40}$ | -2.31 | R3HDM1, UBXN4,<br>LCT, MCM6, DARS1,<br>CXCR4, THSD7B |
| GBR | 12 | 1.11 | 1.13 | $3 \times 10^{-18}$ | -1.50 | ATXN2, BRAP,<br>ACAD10, ALDH2,<br>MAPKAPK5,<br>TMEM116, ERP29,<br>NAA25, TRAFD1,<br>RPL6, PTPN11,<br>RPH3A |
| BEB | 16 | 0.34 | 0.47 | $3 \times 10^{-17}$ | -1.45 | SHCBP1, VPS35,<br>ORC6, MYLK3,<br>C16orf87, GPT2 |
| ESN | 19 | 0.43 | 0.44 | $4 \times 10^{-17}$ | -2.19 | PSG5, PSG4, PSG9,<br>CD177, TEX101,<br>LYPD3, PHLDB3,<br>ETHE1, ZNF575,<br>XRCC1 |
| ESN | 6 | 0.53 | 0.53 | $5 \times 10^{-16}$ | -2.74 | TMEM14A |
| JPT | 1 | 0.75 | 0.76 | $4 \times 10^{-15}$ | -2.27 | ACADM, RABGGTB,<br>MSH4 |
| FIN | 15 | 0.28 | 0.29 | $4 \times 10^{-15}$ | -1.73 | GOLGA8F, GOLGA8G |
| GWD | 6 | 0.59 | 0.62 | $9 \times 10^{-14}$ | -2.25 | KHDRBS2 |
| KHV | 2 | 0.17 | 0.18 | $4 \times 10^{-13}$ | -2.39 | RAD51AP2, VSNL1 |
| TSI | 11 | 0.89 | 0.89 | $5 \times 10^{-13}$ | -1.63 | TYR |

**Table S9:** Most significant balancing selection hits for 1000 Genomes Project Popultions

| Population | Chromosome | start | end | p value | Average $\hat{\beta}$ | genes |
| --- | --- | --- | --- | --- | --- | --- |
| ASW | 6 | 0.322 | 0.327 | $< 10^{-38}$ | 1.33 | RNF5, AGER, PBX2, GPSM3, NOTCH4, TSBP1, BTNL2, HLA-DRA, HLA-DRB5, HLA-DRB1, HLA-DQA1, HLA-DQB1, HLA-DQA2 |
| YRI | 4 | 0.989 | 0.991 | $< 10^{-38}$ | 4.71 | METAP1 |
| IBS | 1 | 1.529 | 1.532 | $< 10^{-38}$ | 6.43 | SPRR4, SPRR1A, SPRR3, SPRR1B, SPRR2D, SPRR2A, SPRR2B, SPRR2E, SPRR2F, SPRR2G |
| ESN | 4 | 0.854 | 0.855 | $< 10^{-38}$ | 5.78 | ARHGAP24 |
| ASW | 5 | 1.355 | 1.355 | $< 10^{-38}$ | 6.61 | NEUROG1 |
| ACB | 11 | 0.051 | 0.051 | $< 10^{-38}$ | 5.16 | OR52E1 |
| ASW | 6 | 0.330 | 0.331 | $< 10^{-38}$ | 4.98 | HLA-DPA1, HLA-DPB1 |
| IBS | 13 | 0.368 | 0.368 | $< 10^{-38}$ | 6.59 | RFXAP |
| STU | 1 | 2.315 | 2.315 | $< 10^{-38}$ | 5.87 | TSNAX |
| STU | 20 | 0.235 | 0.236 | $< 10^{-38}$ | 5.76 | CST9L |

## Footnotes

1 One standard practice is to subtract the genomewide mean of the test statistic from local estimates. But this assumes that the bulk of the genome is evolving neutrally, and recent work has questioned the validity of this assumption (McVicker et al., 2009; Cai et al., 2009; Lohmueller et al., 2011).

2 A third type of selection, background selection, alters genetic diversity in a way that is indistinguishable from shrinking the effective population size (Charlesworth, Morgan, and Charlesworth, 1993), and is therefore not captured by our approach.

3 This model should not be confused with the *β*-coalescent (Schweinsberg, 2003), which is a more general type of coalescent model that allows for multiple merger events. We discuss possible connections between generalized coalescent processes and our model in Section 4.

